# Confrontation of AlphaFold2 models with cryo-EM and crystal structures enlightens alternate geometries of the CYP102A1 multidomain protein

**DOI:** 10.1101/2022.03.21.485149

**Authors:** Philippe Urban, Denis Pompon

## Abstract

Large range structural dynamics plays a critical role for the function of electron transfer proteins. This information is generally not available from crystallographic structures, while cryo-electron microscopy (cryo-EM) can provide some elements but frequently with a degraded spatial resolution. Recently, AlphaFold-based structural modelling was extended to the prediction of protein complexes. In this work, bacterial CYP102A1 from *Priestia megaterium* was used as a test case to evaluate the capability of AlphaFold2 to predict alternative structures critical for catalysis. CYP102A1 monooxygenase, a NADPH-supported fatty acid hydroxylase, works as a soluble homodimer, each monomer harboring two flavins (FAD and FMN) and one heme cofactors. Large conformational changes are required during catalytic cycle to allow successive electron transfers from FAD to FMN and finally heme iron. We used the recently released AlphaFold2_advanced notebook (AF2A), to predict the possible alternate conformations supporting electron transfers in CYP102A1 homodimer. Challenging AF2A-derived models with previously reported experimental data revealed an unforeseen domain connectivity of the diflavin reductase part of the enzyme. Intermolecular crossed complex constitutes a novel type of structural organization never previously described. The predicted formation within the dimer of a stable complex between the heme containing domains was challenged and found consistent with uninterpreted features of reported crystallographic structures and cryo-EM imaging. The particularly efficient CYP102A1 catalytic mechanism was revisited to the light of the new evidenced connectivity in which the FMN-binding domain of each monomer oscillates on themselves to alternatively receive and transfer electrons without needing large structural change in the dimer. Such model was found explanatory for previously contradictory reported biochemical data. Possibility to mimic CYP102A1 structural organization into bicomponent eukaryotic P450 systems was evaluated by designing and modeling in silico synthetic reductase domains built from composite sequence segments from *P. megaterium* and human origins. More generally, this work illustrates how the ability of AF2A to predict alternate complex structures can enlighten and explain conformational changes critical for bio-assemblies.

## Introduction

AlphaFold2-based modeling of protein structures is now recognized as significantly surpassing the performances of previously described computational methods, frequently approaching or potentially overperforming the accuracy of experimental structures^1-3^. Recently, AlphaFold2 capabilities were extended to the prediction of the protein complex structures with a demonstrated efficiency as compared to that of classical docking approaches. This made possible the exhaustive structural predictions for the full proteome of several model organisms^4^ and, in combination with RoseTTAFold, of the protein interactome in *Saccharomyces cerevisiae*^5^. A recently released development version of this algorithm named AlphaFold2_advanced (AF2A) is an interesting alternative to the more finalized “multimer” version^6^, being sometime slightly less precise but much faster, allowing easier user-defined tunings and generating for some complex a range of alternate possible structures.

For many biological systems of interest not only the structure but also the structural dynamics of protein domains within the complex is critical to understand function^7,8^. We asked us whether some poorly documented properties of AF2A could also shed light on these dynamics. To do that we chose as a test case the bacterial CYP102A1 from *Priestia megaterium* (formerly known as *Bacillus megaterium*), a cytochrome P450 (P450) that catalyzes fatty acid hydroxylation^9^. Its polypeptide chain is modular: at the N-terminus a 54 kDa P450 heme-containing domain with its typical heme-thiolate fold^10^ followed by a 65 kDA diflavin reductase domain (the reductase domain) composed of an FMN-binding domain, and a FAD-binding domain. The structure organization of CYP102A1 reductase domain is typical of that of eukaryotic NADPH-cytochrome P450 oxidoreductases (CPR)^11,12^, assimilatory sulfite reductases (SiR) ^13^, methionine synthase reductase^14^, and nitric oxide synthases^15^, which do not include heme domain fusion.

The unusual homodimer structure of the catalytically competent form of CYP102A1^16^ contrasts with the monomeric form of most P450s. Its heme containing domain is also covalently fused to a reductase domain, which is normally present as separate and monomeric chain in most of bacterial and eukaryotic monooxygenase P450 systems that exist as soluble (in prokaryotes) or membrane-bound (in eukaryotes) forms. Multiple reports based on CYP102A1 site-directed mutants and their biochemical analysis led to the dominant opinion that the electron transfers involved exclusively intermonomer mechanisms facilitated by the dimeric structure^16,17^. In addition, extensive analysis of chemical reticulation patterns by mass spectrometry brought some light on the intermonomer organization by identifying putative interchain contacts^18^. Several partial CYP102A1 crystallographic structures, including that of the isolated P450 domain (P450d)^19^ and that of the isolated FAD-binding domain (FADd)^20^ were reported. A fragment of CYP102A1 encompassing P450d and the FMN-binding domain (FMNd) was also crystallized and the structure of the resulting intrachain complex resolved^21^. In contrast, no crystal structures for the full-length protein, for the isolated reductase domain, or for any domain association in the dimer are available up to now.

More recently, cryo-EM structural imaging allowed to visualize the dimeric structure of the full-length enzyme and was used in combination with available partial crystallographic structures to model the full-length dimer^22^. Low resolution imaging of a simple mutant and higher resolution imaging of a synthetic variant featuring a shortened P450d-FMNd linker allowed acquisition of cryo-EM data at a sufficient resolution to satisfyingly embed the previous crystallographic structures into EM density maps. However, parts of the too mobile linker regions were not visible on crystal structures or difficult to attribute in the EM density map and were arbitrary modeled as polyalanine stretches. This resulted in a proposed model where FMNd and FADd of the same monomer forms an intramolecular complex, while FMNd of one monomer is contacting the P450d of the other monomer. Su *et al*. proposed that in a CYP102A1 dimer, the FMN cofactor was receiving electrons from the FAD of the same monomer and transferring its electron to P450d of the other monomer^22^. This model in which the interflavin electron transfer takes place within a closed conformation of CYP102A reductase domain mimics the regular mechanism documented in all eukaryotic CPRs. In such structure, the FAD and FMN xylene rings stands face to face in a close proximity facilitating electron transfers^23^.

This cryo-EM model offered the first experimental view of the full CYP102A1 dimeric structure in a conformation compatible with interflavin electron transfers. Noticeably, cryo-EM data also evidenced the presence of several subpopulations of more opened structures that could constitute alternate conformations. Unfortunately, the corresponding images could not be obtained at a sufficient resolution to raise additional hypothesis on the mechanisms underlying the exceptionally high catalytic efficiency of CYP102A1. The model remained rather obscure concerning the structural determinant permitting the fast FMN to heme electron transfer required to support enzyme activity^24,25^. In the present paper a modelling approach based on AlphaFold2 tools was used to revisit the dimer structure of CYP102A1 and the dynamics supporting the particularly efficient electron transfers involved in its mechanism with a focus on the role of the dimer structure.

## Results and Discussion

### 1. Static and dynamic features of AlphaFold2 modeling

The first version, AlphaFold1, involved a deep-learning and convolution-based approach in which a set of separately PDB-trained modules were used to produce guides, which, combined to physics-based criteria like energy potentials, resulted in predicted models^26^. This approach was improved in the following version, AlphaFold2, by the use of graphs manipulated by attention mechanisms in which coupled sub-networks generated end-to-end models mostly based on geometrical pattern recognitions. Local physics could optionally be applied as a final AMBER refinement step for energy minimization, particularly to optimize side chain geometry.

A particularity of the AlphaFold2_advanced notebook (AF2A) is to give easy access to the several alternate structural models automatically generated from a unique set of amino acid sequences used as input. Generated structures generally constitute a set of highly similar conformers except in the case when alternate and mutually exclusive biological complexes can be formed. Multi-domain protein like the CYP102A1 dimer typically enters in such category. The different situations illustrated in supplementary figure S1 were encountered during complex modelling. First, in the absence of formation of a defined complex, interacting polypeptide chains were individually folded but modeled at stochastic relative positions. Such a situation was illustrated for example during the modeling of a pair of isolated FMNd. In contrast, consistent spatial positioning of modeled domains was systematically predicted when a defined complex can be formed. This was illustrated for example for a dimer of polypeptide chain encompassing both P450d and FMNd. Minor differences between the alternate structural predictions (RMSD < 1.5 Å) were mostly localized in the less geometrically constrained parts of the models. These highly similar structures can then be either averaged, sorted or refined, when necessary, by energy minimization involving the Amber option. A third situation occurred when alternate protein complexes can be formed involving significantly different geometries. In this case, and when confronted to conflictual modeling, alternate instantiations or differences in arbitrations by the AF2A self-attention algorithms resulted in prediction of markedly different conformations from run to run. As recently reported, the reference sequence sets used or the choice of alternate cluster centroid definitions during the multiple sequence alignment step might also play a role in the generation of alternate solution^27^. Considering that the algorithm does not explicitly involve energetic calculation but is mostly based on geometric pattern recognition, such arbitrations are not expected to directly mirror relative thermodynamic stability of predicted complexes. However, geometric complementarity and interaction energy at structural domain interfaces are linked parameters. Such situation clearly took place during AF2A modeling of CYP102A1 homodimer, where at least four alternate geometries can be considered for the association of the three redox domains (see below).

### 2. Prediction of CYP102A1 dimer structure and alternate conformations

The potential of AF2A for *ab initio* prediction of CYP102A1 structure and of its alternate conformations was evaluated. The polypeptide chain of CYP102A1 is 1,049 amino acid residues long, and is a natural fusion between a N-terminal P450 domain and a C-terminal reductase domain that is related to diflavin reductase protein family^9^. Current technical limitations of AF2A implementation did not allow us to easily model more than a total of ∼1,500 amino acid residues in a single run. CYP102A1 dimer accounts for 2,098 amino acids, making single step modelling technically difficult. However, modelling by parts of this dimer followed by the global assembly of these parts remained possible and was evaluated.

The full-length monomer of CYP102A1 can be easily modeled in a single block and the resulting prediction was already released in the AlphaFold Protein Structure Database (APD at https://alphafold.ebi.ac.uk/ with entry P14779. We revisited this automatically generated model by comparing it with structures resulting from ten non-averaged AF2A predictions. The APD-released P14779 structure and all newly structures that we generated always exhibited the same global fold in which the FMNd is complexed to the P450d, while the FADd remains free to move (Supplementary figure S2). The predicted structures for the FADd, FMNd and P450d taken individually were almost identical (RMSDs < 1.5 Å) both between all the AF2A-generated models, and between any of these structures and the available crystallographic references (PDB P450d, 4kew; P450d-FMNd, 1bvy; FADd, 4dqk). RMSDs between the AF2A models and crystal structures for the P450d-FMNd complex (PDB 1bvy)^21^ or of the FADd (PDB 4dqk)^20^ were respectively 0.88 Å and 1.03 Å (data not shown). It can be noticed that in the 1bvy structure an unwanted proteolysis led in reality to the cleavage of the polypeptide chain linking P450d and FMNd, thus finally forming an intermolecular complex^16^. In all of the AF2A-modeled P450d-FMNd complexes, distances and orientations between the two redox cofactors (heme and FMN) were compatible with efficient electron transfers (Supplementary Figure S3). In contrast, the modeled relative orientations of FADd in respect to the FMNd-P450d complex appeared extremely variable within the limits permitted by the FMNd-FADd linker polypeptide chain flexibility (Supplementary Figure S2). This observation is consistent with the high flexibility of the FMNd-FADd hinge region that makes relative orientation of attached domains poorly defined in the absence of complexation. Therefore, AF2A modeling of the full-length CYP102A1 monomer systematically enlighten the formation of a FMNd-P450d complex, while ignoring the possible formation of the alternate FMNd-FADd complex. In contrast, this last complex was always predicted to form during AF2A modeling of the isolated FMNd-FADd part of the enzyme. Consequently, the fold of the CYP102A1 isolated reductase part and the ones found in the closed conformation of eukaryotic CPRs^11^ were predicted to be very similar. However, such finding did not take into account the specific dimeric structures of the bacterial enzyme.

In order to challenge the dimer formation suggested by sedimentation and size-exclusion chromatography data^16,28^, the modeling of a pair of CYP102A1 chains with AF2A was attempted. Considering the computing limitations resulting from the large size of the dimer, a modeling-by-parts strategy was designed taking into account all possible modes of dimer formation (Figure 1). We first focused on a possible autonomous P450d dimerization independently of the reductase contributions. We observed that AF2A systematically (in 10 out of 10 runs) predicted that a pair of isolated P450d can form a well-defined complex (Figure 2). Presence of two P450d chains was also described in the asymmetric units of crystal structures PDB 6h1s and 4kew, but the relative P450d-P450d geometries in the two crystals were markedly different and differed from the geometry predicted by AF2A. The P450d structures taken individually in the predicted dimer are highly similar to crystal structures from PDB 6h1s and 4kew with a RMSD of 0.77 Å (413 Cα/453) and of 0.74 Å (381 Cα/400], respectively. The free energy for the formation of the P450d-P450d interface was estimated using the Prodigy webserver^29^ to be -9.3 kcal/mol for the AF2A model compared to -5.2 and -7.0 kcal/mol for the associations seen in the asymmetric units of the discussed crystal structures. Dimerization of isolated CYP102A1 P450d was not experimentally observed to occur spontaneously in solution since purified P450d eluted as a monomer in size exclusion chromatography^16^. In contrast, some sedimentation velocity experiments on partially proteolyzed CYP102A1 have repeatedly shown that, besides the homodimer, a ‘1.45-mer’ was also prominent in solution^28^. This unusual stoichiometry was interpreted as being the result of the binding of a full-length CYP102A1 monomer to an isolated proteolyzed P450d. These contradictory results evidenced that a pair of isolated CYP102A1 P450d domains can form a defined complex or not, depending on experimental conditions. More recently, cryo-EM reports illustrated that in the dimeric form of the full-length CYP102A1, the two P450d subparts are associated into a compact structure^22^. However, it was unclear whether such contact resulted from direct heme domain interaction or from some geometric constraints mediated by the attached reductase parts.

**Figure 1.**
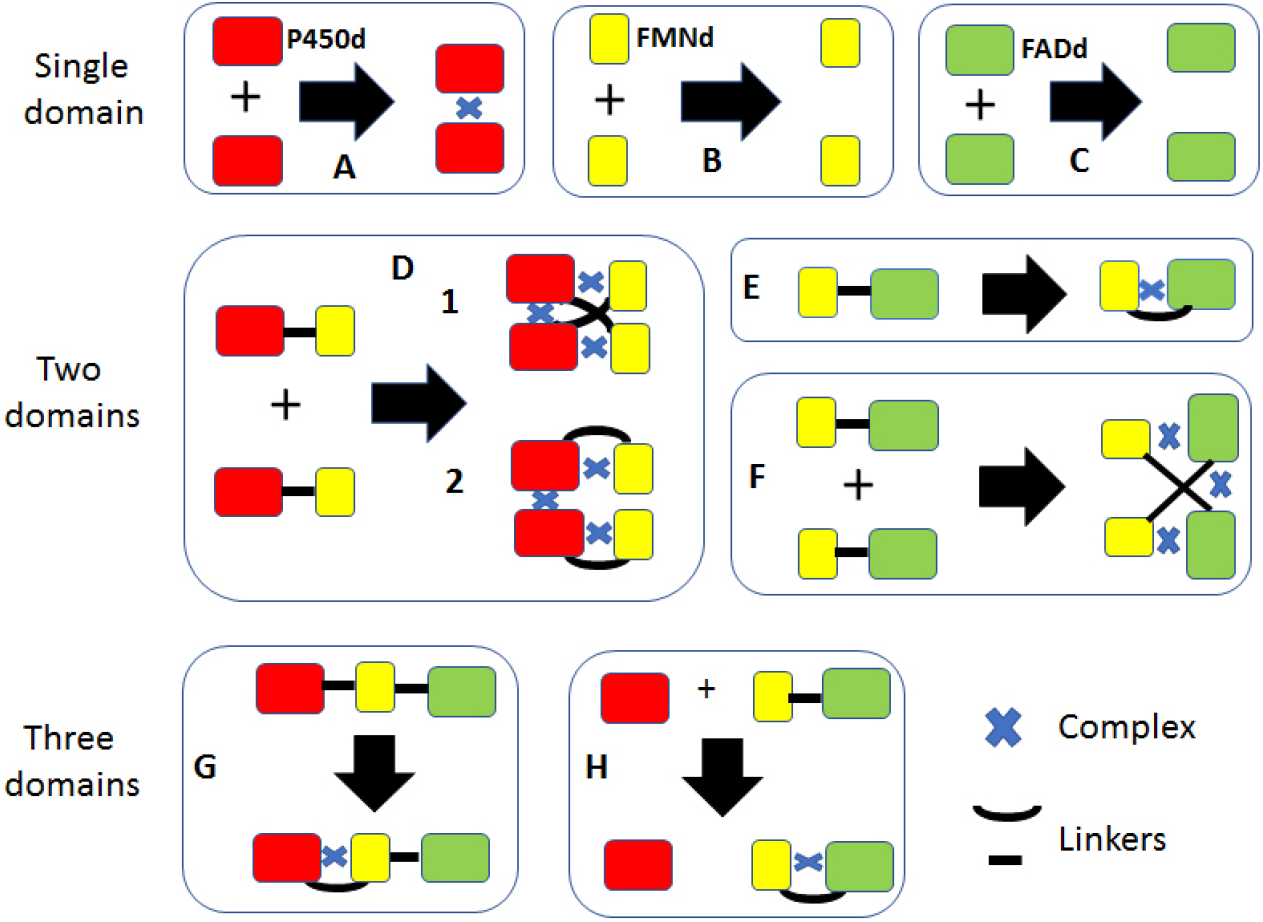
Description of the strategy followed for the AF2A modelling by parts to establish the alternate structures of full length CYP102A1 dimers. P450d, FMNd and FADd are represented as red, yellow and green boxes, respectively. Polypeptide chain linkers considered during modelling process are indicated by solid lines and the predicted complex formations with a blue cross. Five to ten AF2A modeling attempts were performed per case. *Panel A*. A pair of P450d chains formed a dimeric structure. *Panels B and C*. A pair of FMNd or FADd chains never formed a complex. *Panel D*. A pair of P450d-FMNd chains formed a complex either in a *trans*-configuration (D1), or in a *cis*-configuration (D2). *Panel E*. A FMNd-FADd monomer folded in a closed conformation in which FMNd and FADd are complexed. Panel F. A pair of FMNd-FADd formed a crossed complex in which the FMNd of one chain was complexed to the FADd of the other. *Panel G*. The monomer of full-length CYP102A was modeled in a structure in which the FMNd is complexed to the P450d. *Panel H*. The presence of P450d did not modify the folding of the FMNd-FADd monomer that remained identical to case E.

**Figure 2.**
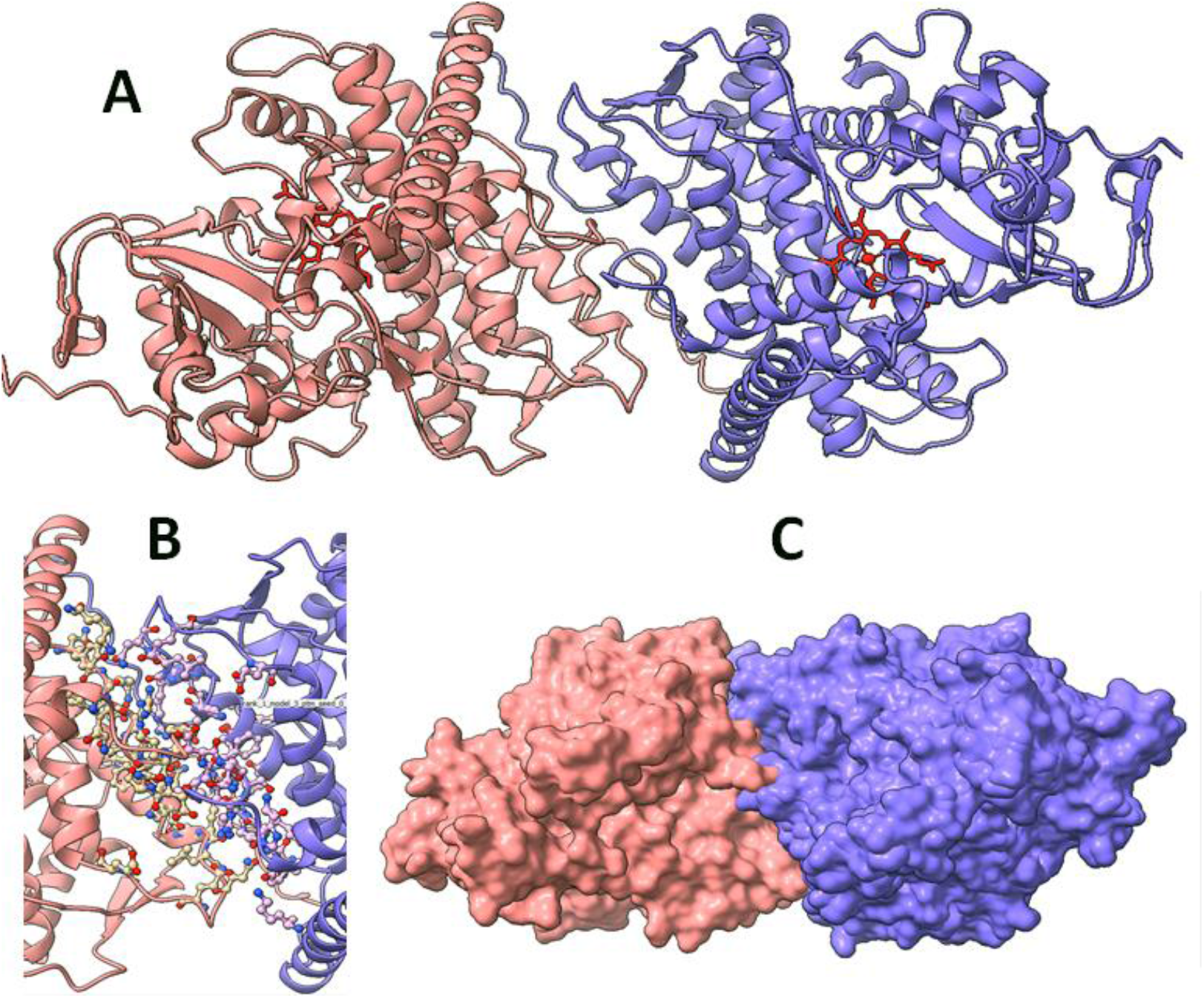
AF2A modelling of dimer formation from a pair of P450d (residues 2-458). *Panel A*. Global view of the predicted P450d-P450d dimer. The two chains are colored differently for the sake of clarity. The two heme cofactors were colored in red. *Panel B*. Enlarged view of the P450d-P450d interface with the side chains of interacting amino acids represented as stick models colored by elements (blue, nitrogen; red, oxygen; and white, carbon atoms). *Panel C*. Surface envelope of the two complexed P450d illustrating the compact structure of the dimer (same coloring as in panel A).

Cryo-EM imaging of the full length CYP102A1 dimer illustrated that the two reductase domains are associated to each other by a FADd-FADd interface expected to contribute to dimer stability. The FMNd-FADd complex was modeled into EM density maps in a way mimicking the closed conformation found in all eukaryotic CPRs. Consequently, formation of the resulting intra-molecular complexes was not expected to contribute to dimer stabilization. To challenge this view, the structure of a pair of reductase domains (lacking P450d) was AF2A modeled. Surprisingly enough, this resulted into a model sharply contrasting with previous cryo-EM interpretations (Figure 3). In the predicted structure, the FMNd of one monomer formed a complex with the FADd of the other monomer and reciprocally. Such a crossed geometry was never described for any diflavin reductases. While the AF2A-predicted chain connectivity drastically differed from the cryo-EM model, the global shape of the reductase dimer remained very similar and consistent with EM imaging as discussed below.

**Figure 3.**
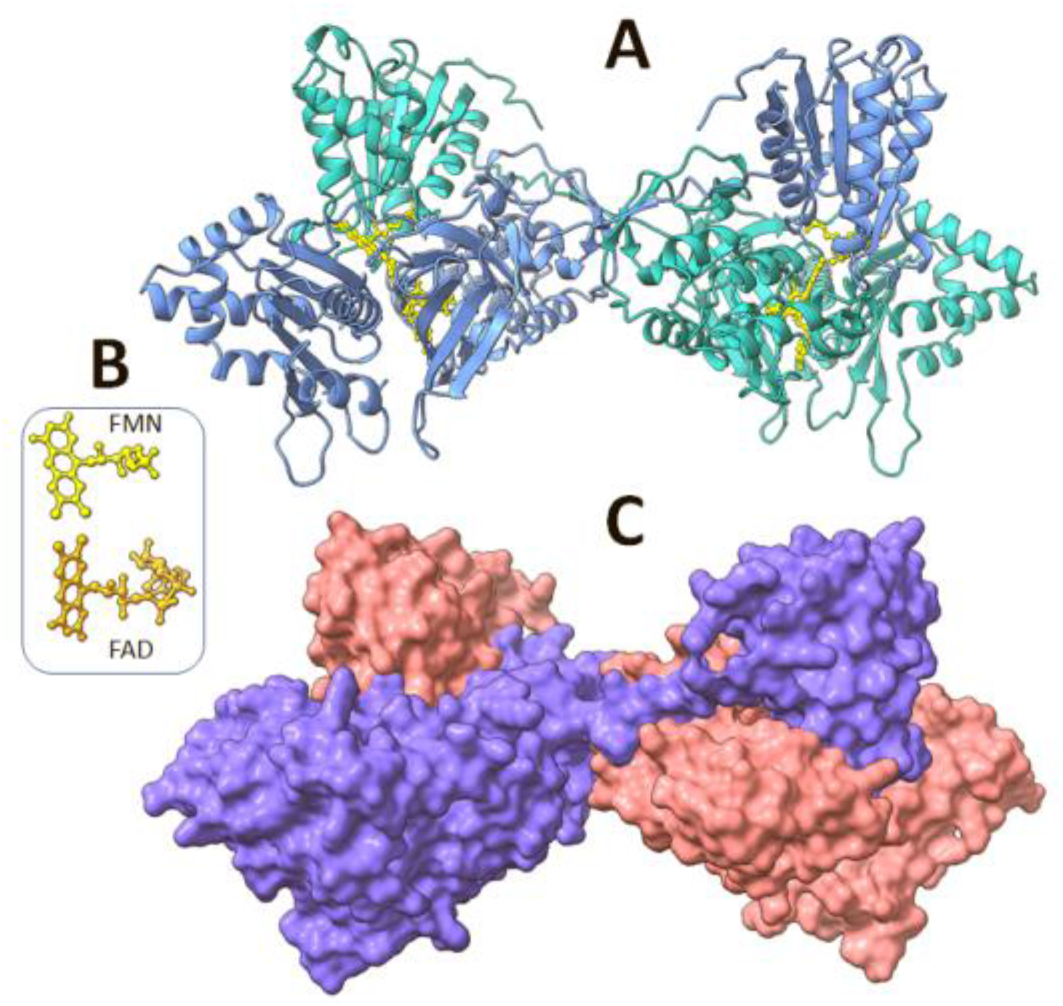
AF2A predicted crossed dimer formed from a pair of FMNd-FADd. *Panel A*. Structure of the dimer in which the FMNd of one chain (seen at top) forms a complex with the FADd of the other chain (seen at bottom). One chain is colored turquoise, the second is in blue. The four flavin cofactors (two FADs and two FMNs per dimer) are shown in stick models colored yellow. They were placed within the AF2A-predicted model by overlay and RMSD minimization of the crystal structures of FMNd (in PDB 1bvy) and of FADd (PDB 4dqk). *Panel B*. Relative orientation of FMN and FAD cofactors extracted panel A, showing the close location of both cofactors facing each other by the xylene rings of their isoalloxazine moiety. *Panel C*. Surface envelope of model from panel A.

In our model, dimer formation resulted from the three different types of interactions as illustrated in Figure 4. Binding free energy (ΔG) resulting from the inter-chain complex formation between one FMNd and one FADd was estimated to be -10.5 kcal/mol, thus contributing for -21 kcal/mol to the dimer stability considering the symmetry. The -10.5 kcal/mol value was very similar to the -9.9 kcal/mol value estimated for the corresponding intra-chain complex in the monomeric reductase structure. In the dimer, the interaction between the FADd of one monomer and the linker region connecting FMNd to FADd of the other monomer additionally contributed for 2 x -3.6 kcal/mol (due to symmetry), and the interface between the two FADd for -4.9 kcal/mol. The total binding energy between the two reductase chains was thus estimated to be ∼ 33 kcal/mol in the crossed dimer compared to -5 kcal/mol mediated by the unique FADd-FADd interface in the cryo-EM model interpretation. Such value of 5 kcal/mol seems to be hardly consistent with the formation of a stable complex and the evidenced experimental affinity for CYP102A1 dimerization in the nM range^16^. Consistently, no complex formation was predicted during AF2A modeling of a pair of isolated FADd. Moreover, we also modeled pairs of three eukaryotic CPRs (human, yeast and *A. thaliana* ATR2), but none of them was predicted to form a dimer, consistently with their known monomeric structures. The formation of a crossed complex thus appeared to be a specific feature stabilizing the dimeric form of CYP102A1.

**Figure 4:**
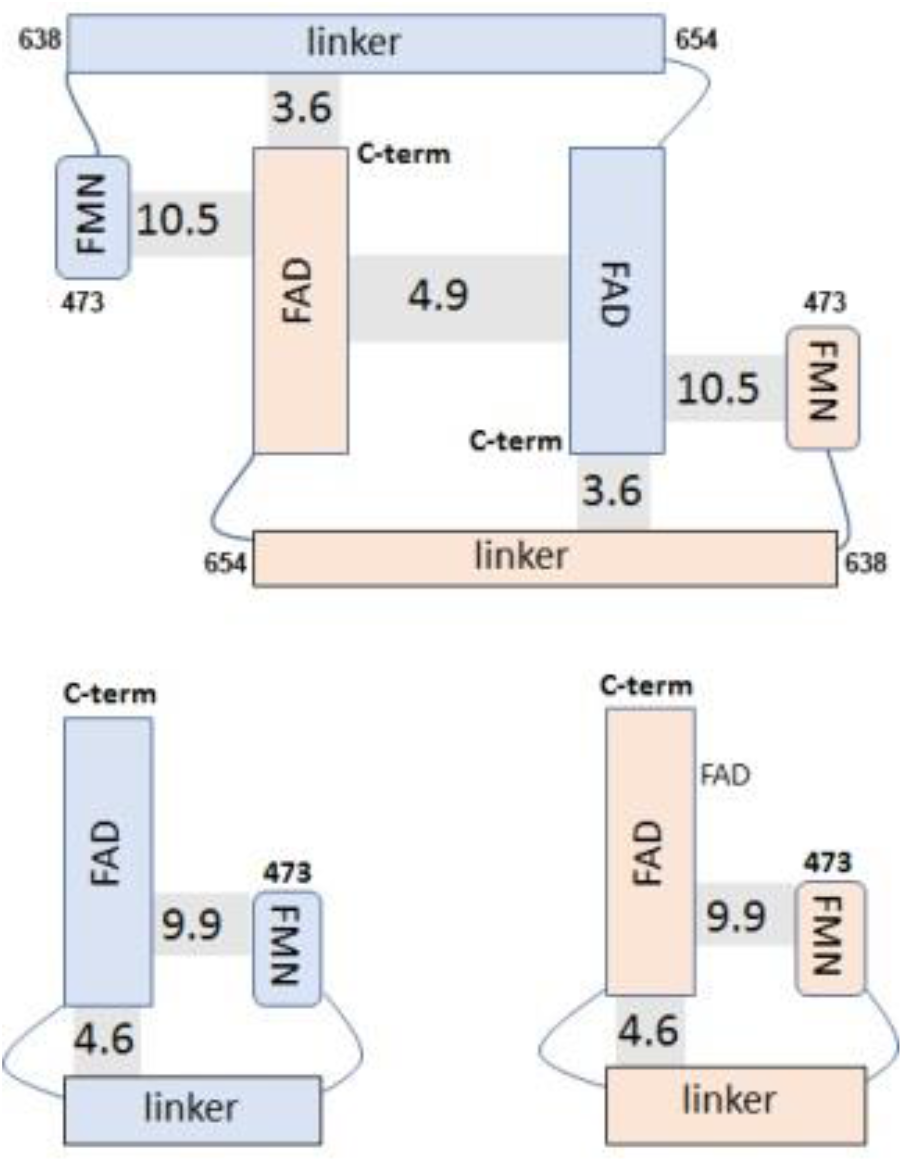
Comparison of interchain and intrachain free energies (ΔG) contributing to structural interactions in the case of the FMNd-FADd dimer (top) and monomer (bottom). Free energies were calculated using the Prodigy algorithm on AF2A predicted structures. The limits of the different considered domains used during calculations are indicated by the CYP102A1 sequence positions. For calculations, the global structures were split at the indicated position keeping atom coordinates unchanged. Energies are given as kcal/mol in the shaded area of each interdomain region.

The next step was to predict the relative structural organization of the P450 and reductase domains in the dimers. To do so, we first AF2A modeled two CYP102A1 fragments each encompassing P450d and FMNd without FADd. The resulting AF2A-predicted structure was found to be dimeric and to involve a P450d-P450d interface identical to the one found in the dimer of isolated P450d fragment in PDB 1bvy (See Figure 2 and supplementary figure S4). In the P450d-FMNd dimer, the two FMNd formed symmetrical complexes with the proximal faces of P450s opposite to the P450d-P450d interface. The configuration of the P450d-FMNd complex was found to be similar to the corresponding complex predicted for the full-length CYP102A1 monomer, and also to the one observed in the crystal structure PDB 1bvy (RMSD of 0.77 Å). Such dimeric complex thus preserves the structural organization suitable for efficient FMN to heme electron transfer (Supplementary Figures S3). Considering now the polypeptide chain topology, the polypeptide linkage between P450d and FMNd can involve either a *cis*-or a *trans*-configuration of the dimer. In the first case, an intra-chain electron transfer would occur, when the *trans* configuration would only permit inter-chain electron transfer. Interestingly, repeated AFA2 runs stochastically generated the two alternate geometries of the P450d-FMNd dimer with a similar frequency when modeling a pair of a deletion variant with a 12-residue shortened P450d-FMNd linker (Δ^457^GGIPSPSTEQSA^468^). The truncated variant was exploited in the cryo-EM study to significantly improve map resolution while mostly causing P450 function inactivation^22^. On the other hand, when modeling a pair of the wild-type sequences we exclusively obtained the dimer in *cis*-conformation. AF2A models also revealed that geometry of the P450d-FMNd complex has tendency to be modified with the variant sequence by tilting the FMNd relative to P450d in a way reducing stretch on the shortened linker. It could be hypothesized that the inactivation of the deletion variant might be caused by this FMNd tilting, thus impairing electron transfer. The difference of AF2A-predicted frequencies for the *cis*- and *trans*-configurations between the deletion variant and wild type enzyme may result from the higher geometrical constraints to be supported by the shortened linker in the *cis*-configuration. However, FMNd must alternatively form complexes with P450d and FADd during catalytic cycles to alternatively reduce and oxidize its FMN cofactor. In dimer, such conformational changes are most likely asynchronous between the two monomers leading to mixed geometries potentially releasing constraints that very likely exist in symmetrical structures. It is thus difficult to conclude whether the P450d-FMNd *trans*-configuration interpretation reported in the cryo-EM study, based on experiments with a mostly inactive variant, was or not representative of the wild type geometry.

Two complementary mechanisms could thus contribute to dimerization of the full length CYP102A1. The first one relates to the autonomous dimerization of the P450 domains, the second results from the formation of a symmetrical FMNd-FADd crossed complex. A significant difference between these two mechanisms is that P450d homodimer does not need to be broken to permit catalytic cycle, while the crossed reductase complex must dynamically rearrange to permit the alternate FMNd complexations with FADd and P450d. However, CYP102A1 dimerization can be maintained during catalytic cycles by both the P450d-P450d and the FADd-FADd interfaces, not accounting for the asynchronous disruption of FMNd-FADd complexes during the FMNd catalytic flip-flops.

### 3. Assembly of full-length CYP102A1 dimer model with the reductase in a closed conformation

A composite model of the full length CYP102A1 dimer featuring a closed conformation of the diflavin reductase part was built by combining AF2A modeling for the P450d dimer and for the FMNd-FADd crossed dimer (See Methods Section). Being a homodimer of two identical CYP102A1 chains, the two partial structures must share a common symmetry axis within the final global model. Individual symmetry axis for the two modeled parts were thus calculated and suitable translation and rotation applied to coordinates so as to match the symmetry axes of partial models. Once aligned, only two parameters have to be adjusted to build the final model: the distance between the barycenter’s of partial models and a rotation angle along the common symmetry axis. Distance was adjusted to the shorter value not creating structural clashes between the two modeled parts and the unique angle parameter was selected by optimizing the embedding quality of resulting model into the previously reported cryo-EM density maps^22^. AF2A generated models were worked as rigid bodies during all the steps.

The resulting full-length CYP102A1 model (Figure 5) exhibits a closed reductase conformation and was found to tightly fit into reported EM density maps as illustrated under different view angles on Figure 6. Simulated electron densities for predicted structures appeared highly consistent with experimental cryo-EM density maps at the levels of P450d and FADd models with a similar RMSD of 2.4 Å. However, the part of the cryo-EM density around FMNd appeared significantly larger than the maximum volume that can be occupied by this domain (red-framed in Fig.6A). Such feature suggested that this enlarged EM map could result from the averaging of densities resulting of alternate FMNd conformations. This hypothesis makes sense considering that the FMNd must form alternate complexes with FADd and P450d to support electron transfers required for enzyme function. Further adjustment of the FMNd position in the model was not attempted because it would disrupt the geometry required for the electron transfer with FADd and will not help in better fitting a too large map. One other minor misfit illustrated in Figure 6 is the absence of predicted AF2A structure compared to experimental EM maps close to the N-termini of the two monomers (red-framed in Fig. 6D). This can be easily explained by the absence in the AF2A modeling of the His-tag extension present at the N-termini of the proteins used for cryo-EM imaging.

**Figure 5.**
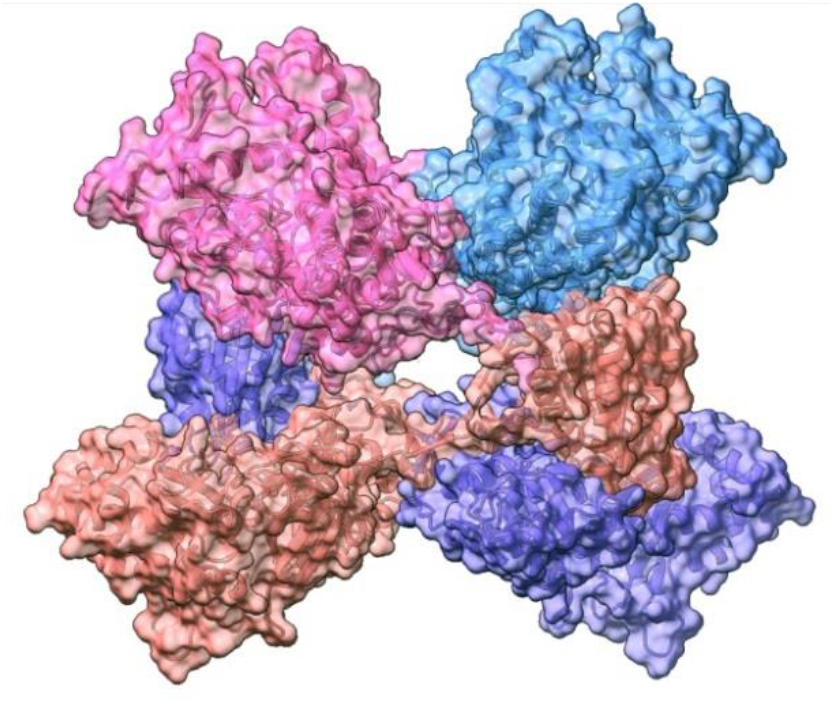
Model for the full-length CYP102A1 dimer with the reductase domains in the closed conformation. The model was assembled from the AF2A predicted structures of the P450d dimer and of the FMNd-FADd dimer as presented in the text. P450d stands at top, and the reductase domains at the bottom. The orange-colored reductase domain (FADd and FMNd) and the pink-colored P450d belong to the same monomer. The blue (P450d) and violet-colored (reductase) domains belong to the other monomer. In contrast, in the noncrossed P450dFMNd configure-tion, the orange-colored FMNd and the pink P450d would be associated together.

**Figure 6.**
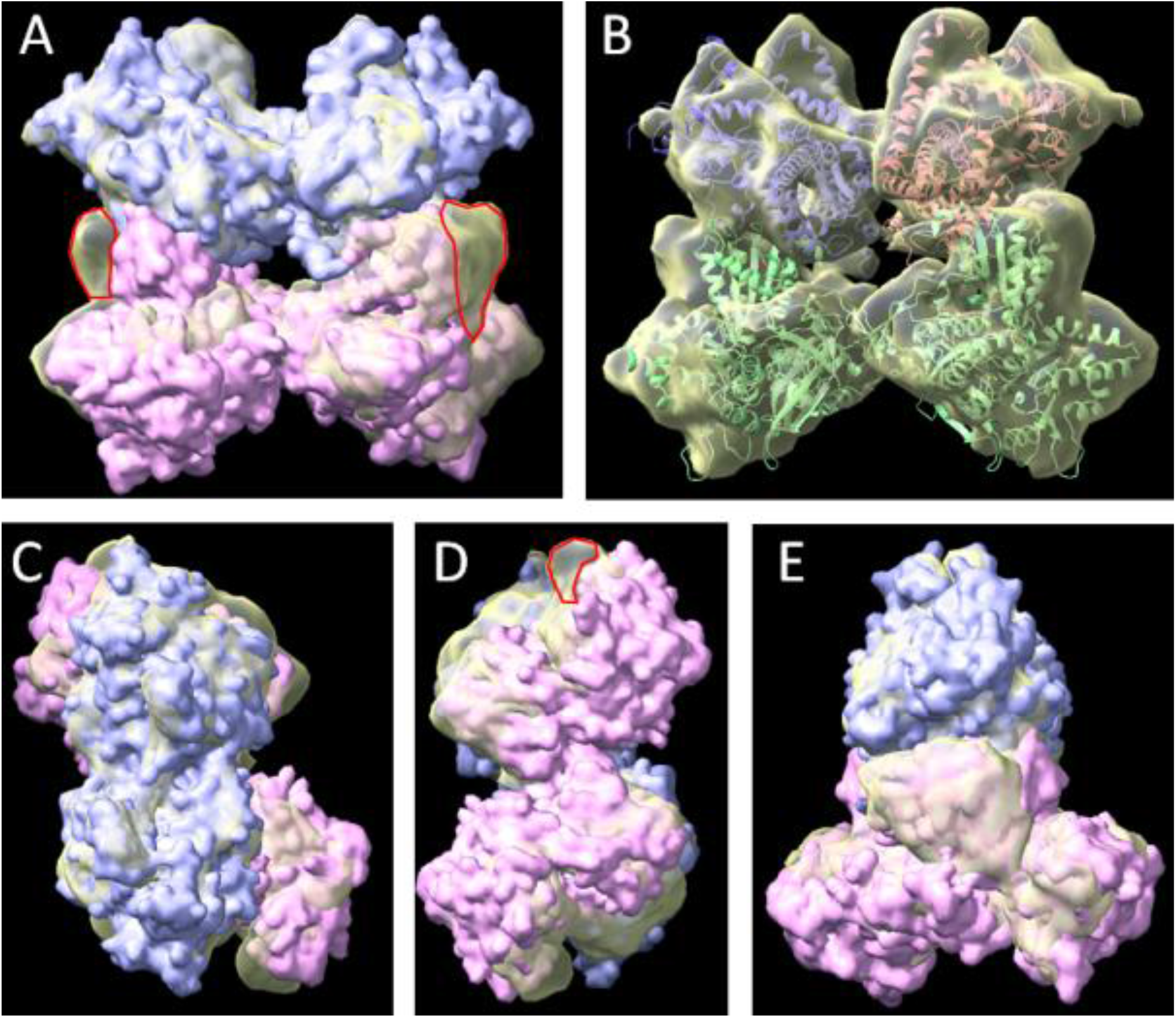
Comparisons of AF2A-predicted structures and experimental cryo-EM envelope of full-length CYP102A1 dimer with the reductase domains in a closed conformation. The AF2A model and its theoretical electron density map were embedded into the experimental EMD-20785 density map as described in the Methods section. *Panels A, C, D, and E* represent the same structure seen from different angles (face, top, bottom and side, respectively). Electron density for the predicted structures was modeled using the ChimeraX^30^ molmap plugin assuming a 4 Å resolution. Calculated density for P450d-P450d dimer (blue) and FMNd-FADd dimer (pink) were overlay with the cryo-EM EMD-20785 map (yellow) at its experimental 7.6-Å resolution. *Panel A*. The red framed area of the cryo-EM envelope highlights the experimental density that was not filled by electron density of the FMNd in the closed conformation. *Panel B*. Same models in ribbon style embedded into the cryo-EM density map.

The P450d-P450d interfaces resulting from the cryo-EM and the AF2A approaches were compared in more details (Figure 7). As illustrated, the P450d-P450d interface was highly similar between the AF2A predicted and the cryo-EM experimental structures, including for the nature and position of contacting amino acid residues. Structure similarity search in the PDB allowed us to identify several structures including two P450 domains per asymmetric unit, but only one (PDB 1bvy) allowed a tight match between the AF2A model and relative positioning of P450d in the crystal structure (Supplementary Figure S4). Totally different relative geometries were observed for the crystal structures PDB 6h1s and 4kew that also include two P450 domains per asymmetrical unit.

**Figure 7.**
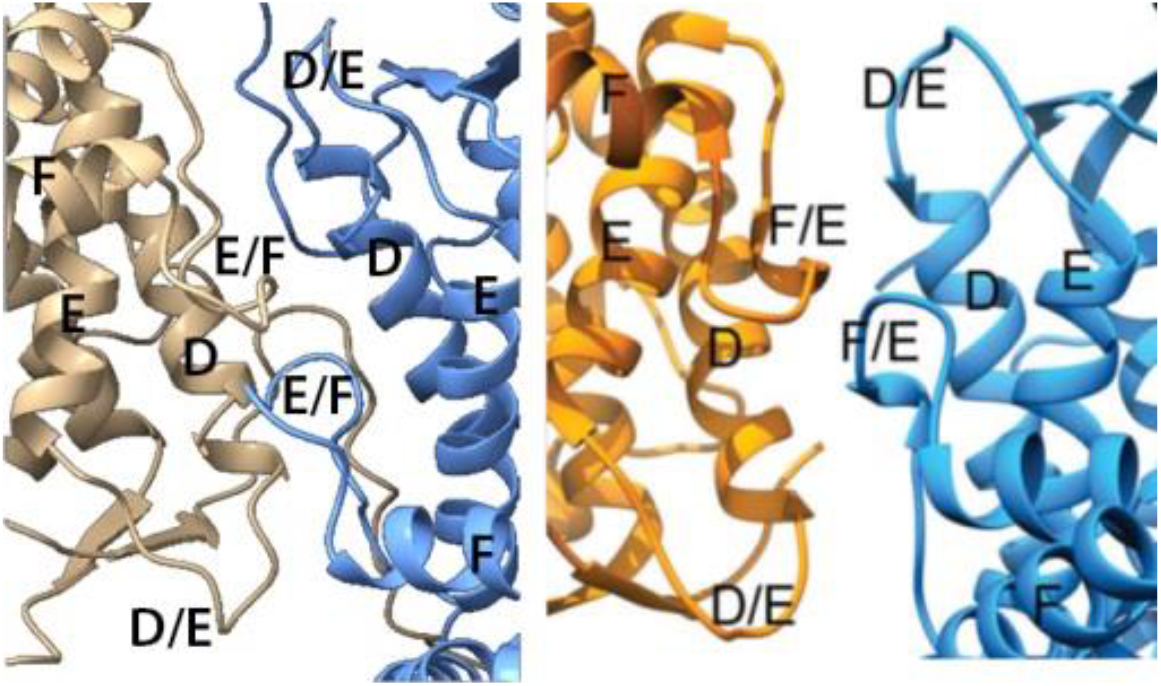
Comparison in similar orientation of the P450d-P450d interface in the AF2A predicted dimer (Left panel) and in the published experimental structure^22^ derived from the docking of crystallographic models into the cryo-EM density map (reproduced from figure 2D^22^). (Right panel). The two P450d in the dimer are colored differently for the sake of clarity. The lettering in each figure corresponds to the P450 helices and loop identifications as given in the published paper.

A major difference between the asymmetrical units of these three crystals is that only structure 1bvy also includes a supplementary FMN-binding domain complexed to one of the two P450d copies^21^. Formation of the P450d-FMNd complex was clearly geometrically incompatible with the crystal packing of the 4kew and 6hs1 structures, thus impairing formation in 1bvy crystal of non-physiological dimeric structures likely resulting from packing constraints. Predicted and cryo-EM determinations of the structure of the dimeric reductase domain were also compared. Taken individually, FMNd and FADd folds were highly similar between AF2A model and corresponding crystal structures used to build the cryo-EM models (Supplementary Table S1). Similarly, the structure of the FADd-FADd interface deduced from cryo-EM data and the corresponding interface in the AF2A model involved the same structural elements and contacting residues (Figure 8). This confirmed the power of AF2A to accurately predict complex interfaces that are not described in experimental full-length structures. However, AF2A and cryo-EM models significantly differed at the level of the polypeptide chain linking FMNd and FADd. In the crossed dimer structure, the FMNd-FADd linker encompasses not only the 12-residue long hinge segment (residues 632-643) but also the N-terminal loop of FADd (residues 644-654) crossing the FADd-FADd interface. This chain segment is crossing two times (one for each monomer) the FADd-FADd interface in the AF2A model of the dimer, thus allowing the formation of the crossed interchain complex. In contrast, no chain was crossing the interface in the cryo-EM model that involves two intramolecular FMNd-FADd complexes. To solve this contradiction, the high resolution cryo-EM density map EMD-21100 was reexamined. The elements of the two FMNd-FADd linkers crossing the FADd-FADd interface in the AF2A model were unambiguously visualized on this experimental map and accurately fitted with the electron densities of cryo-EM map (Figure 9). In contrast, the electron density of the loop linking FMNd to FADd in the intramolecular complex predicted by Su *et al*. is not seen in the cryo-EM map^22^. This observation strongly supports a novel alternate structure of diflavin reductase enzymes, in which the FMN domain of one monomer is associated to the FAD domain of the other monomer, contrasting with the closed monomeric conformation model always evidenced in eukaryotic diflavin reductases.

**Figure 8.**
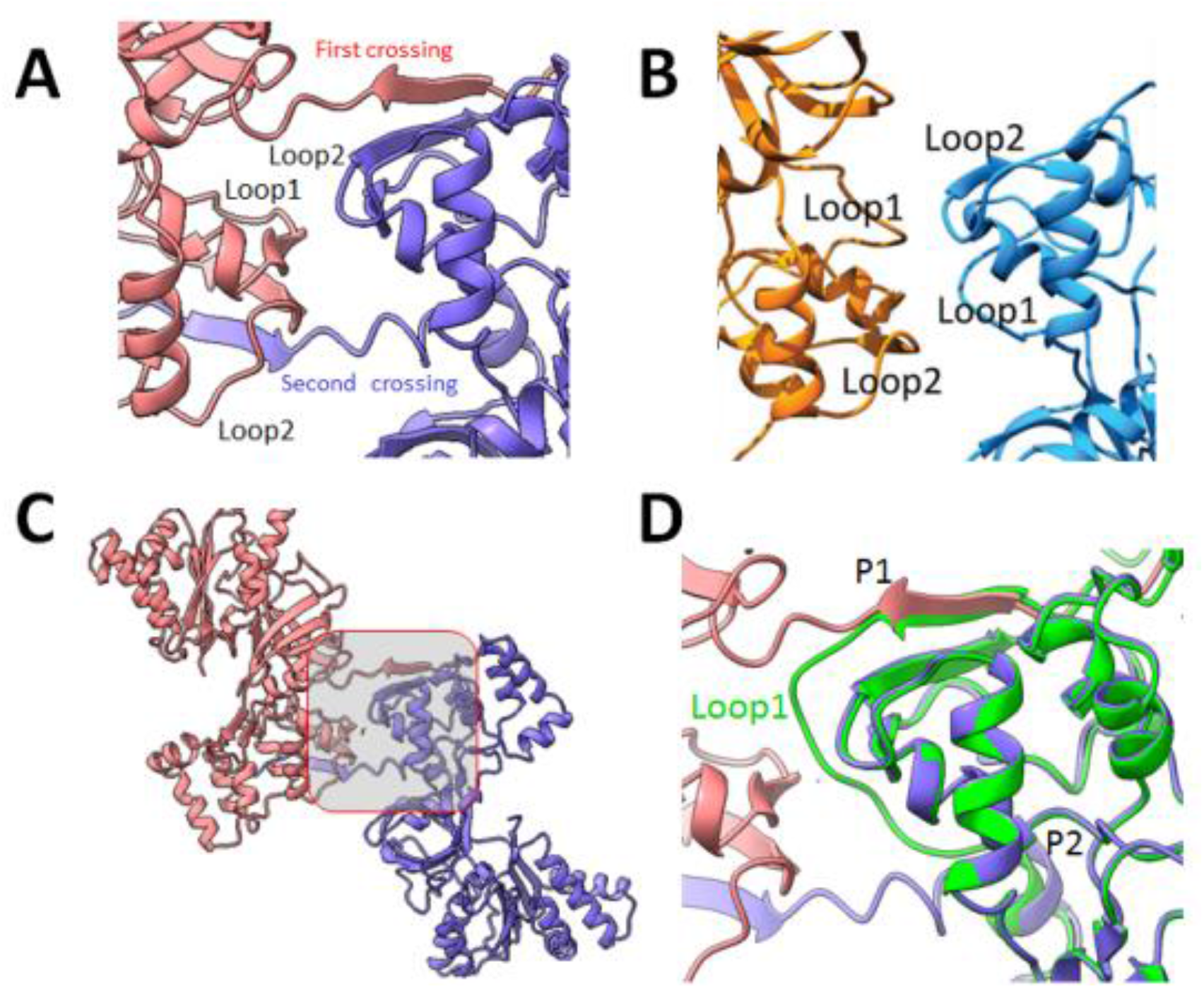
FADd-FADd interface deduced from modeling of a pair of FMNd-FADd chains compared to the reported cryo-EM model. *Panel A*. The interface between the two FADd as seen in the AF2A-predicted structure. Both chains have been differently colored for the sake of clarity. Loop1 and loop2 correspond respectively to the 8-residue loop (from residue 647 to residue 655 in CYP102A1) that cross the interface in the modeled structure. *Panel B*. The interface, in a similar orientation as reported from the published cryo-EM based model (reproduced from figure 2E^22^). *Panel C*, Position of the FADd-FADd interface presented in panel A was underlined into the global model. *Panel D*. Overlay of AF2A-predicted structures of the FMNd-FADd dimer (chains were colored in violet and pink) and of the corresponding monomer (green). The Loop1 exhibits an extended configuration in the dimer crossing the interface while turning back to the FADd of the same chain in the monomer.

**Figure 9.**
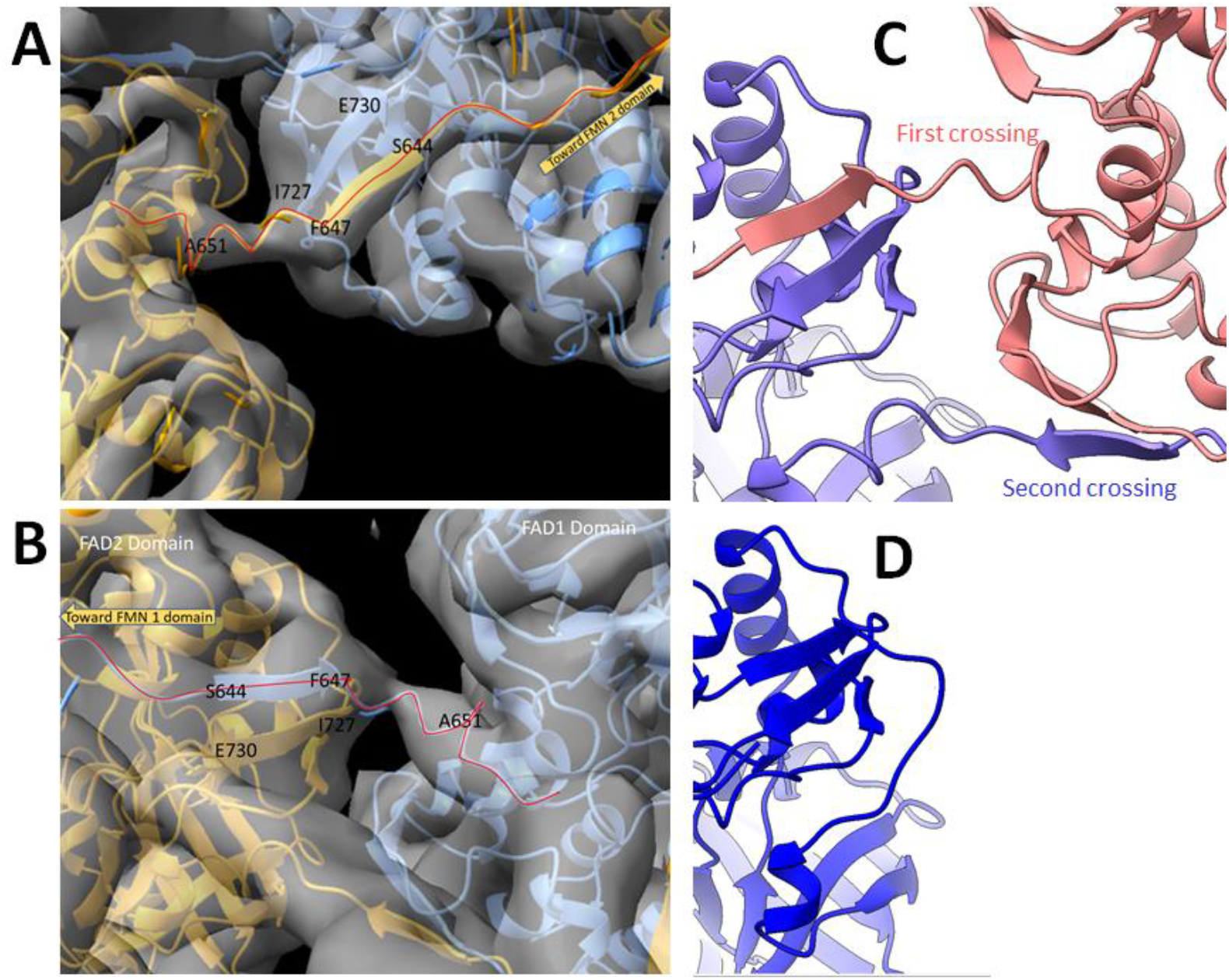
Overlay of the FADd-FADd interface in the AF2A-modeled FMNd-FADd dimer with experimental EMD-21100 density map. *Panels A and B*. The two regions of the cryo-EM map where the AF2A modeled FMNd-FADd linkers are crossing the interface. The red line highlights the course of the crossing chain that links the two flavin domains of the same monomer. The envelope from the cryo-EM density map is shown as a greyish transparent surface overlaying the modeled structure. The polypeptide chains of the dimer in crossed configuration are colored differently for the sake of clarity. *Panel C*. Close view of the FADd-FADd interface showing the two crossing chains in the predicted model of the dimer. *Panel D*. Same view but in the predicted model of the monomer.

Similarly, the amino acid side chain connectivity for the P450d-FMNd linker segment was reexamined. Reported cryo-EM data interpretations favored a crossed geometry based on the fit of the visible electron density using a polyalanine stretch to model the linker in the absence of sufficient resolution permitting to visualize the corresponding region by a more detailed analysis. AF2A modeling predicted that the crossed and the non-crossed geometries are both geometrically possible but difficult to discriminate (Supplementary Figure S5). Reported high resolution cryo-EM maps were experimentally obtained using a 12-residue deletion in the P450d-FMNd linker in order to improve resolution. This deletion that significantly affects the activity of CYP102A1^22^, was also predicted to clearly favor formation of the crossed geometry during AF2A modeling attempts destabilizing the *cis* conformation, as discussed above. The densities attributed to this linker in the cryo-EM maps were thus not necessarily representative of the native structure, letting open the possibility that both crossed and non-crossed geometries could coexist within the native CYP102A1 enzyme.

Confrontation of AF2A modeling to reported experimental data led us to revise previous interpretations based on crystal and cryo-EM data, evidencing that autonomous P450d-P450d dimerization predicted by AF2A was experimentally present but not documented in particular crystal structures and that the reductase domain dimer was mostly stabilized by two chain crossings of the FMNd-FADd linker that encompasses the hinge segment and the N-terminal loop of FADd.

### 4. Alternate assembly of the full-length CYP102A1 dimer model with the reductase in an open conformation and mechanistic consequences

The particularly high turnover of CYP102A1 compared to bi-component P450 systems remained puzzling in regard to the role of the dimerization. Considering the AF2A structures that we have presented and the cryo-EM published structures, the dimer stability resulted from multiple interactions involving P450d-P450d and FADd-FADd interfaces, but also crossed FMNd-FADd complex associations. Successive electron transfers, first from FAD to FMN then from FMN to P450 are required to support a catalytic cycle. Both complexes involve the similar surface area of FMNd. This implies that a given FMNd has to rapidly shuttle between mutually exclusive complexes with FADd and P450d. A dissociation of the dimer followed by successive intra-chain electron transfers can be excluded as requiring a too high energy cost and because the monomer was demonstrated to be inactive^16^. A partial opening of the structure by dissociating at least one of the two FADd-FMNd complexes without disruption of the P450d-P450d and FADd-FADd interfaces would be more consistent. This process would only require a rotation on itself of one of the two FMNd in the dimer, by breaking its interchain FMNd-FADd interface and forming the alternate association with P450d. Such mechanism could be asynchronous between the two monomers and consequently not involve major structural rearrangement. Its feasibility depends on a sufficient length and flexibility of the P450d-FMNd linker and of the FMNd-FADd linker, the latter includes the hinge segment and the long loop at the N-terminus of FADd in the connecting domain.

Contrasting with the closed conformation of the FMNd-FADd complex, the open conformation of the reductase domain allowing formation of the FMNd-P450d complex is shown in Figure 10. This structure of the full-length CYP102A1 was basically generated in the same approach used to get the structure with the reductase in the closed conformation, but by combining the AF2A-predicted structures of the FMNd-P450d dimer and of the isolated FADd dimer. The structure of the isolated FADd dimer cannot be directly AF2A modeled in the absence of its stabilization through formation of the crossed FMNd complexes. Consequently, atomic coordinates specific of the FADd associated in a FADd-FADd dimer were extracted from those in the FADd-FMNd dimer by simply erasing FMNd atoms to avoid domain duplication during full structure assembly. Similarly to the approach used for the global modeling of CYP102A1 closed structure, model embedding into EMD-21099 density map was made by tuning angle around the restored symmetry axis of the two partial structures. Interestingly, the resulting opened reductase model can also be fitted in the EMD-20785 density map that was used to assemble the CYP102A1 closed model, filling the empty space of the FMNd EM envelope that remained empty in the docking of the AF2A closed structure with FMNd in the open structure.

**Figure 10.**
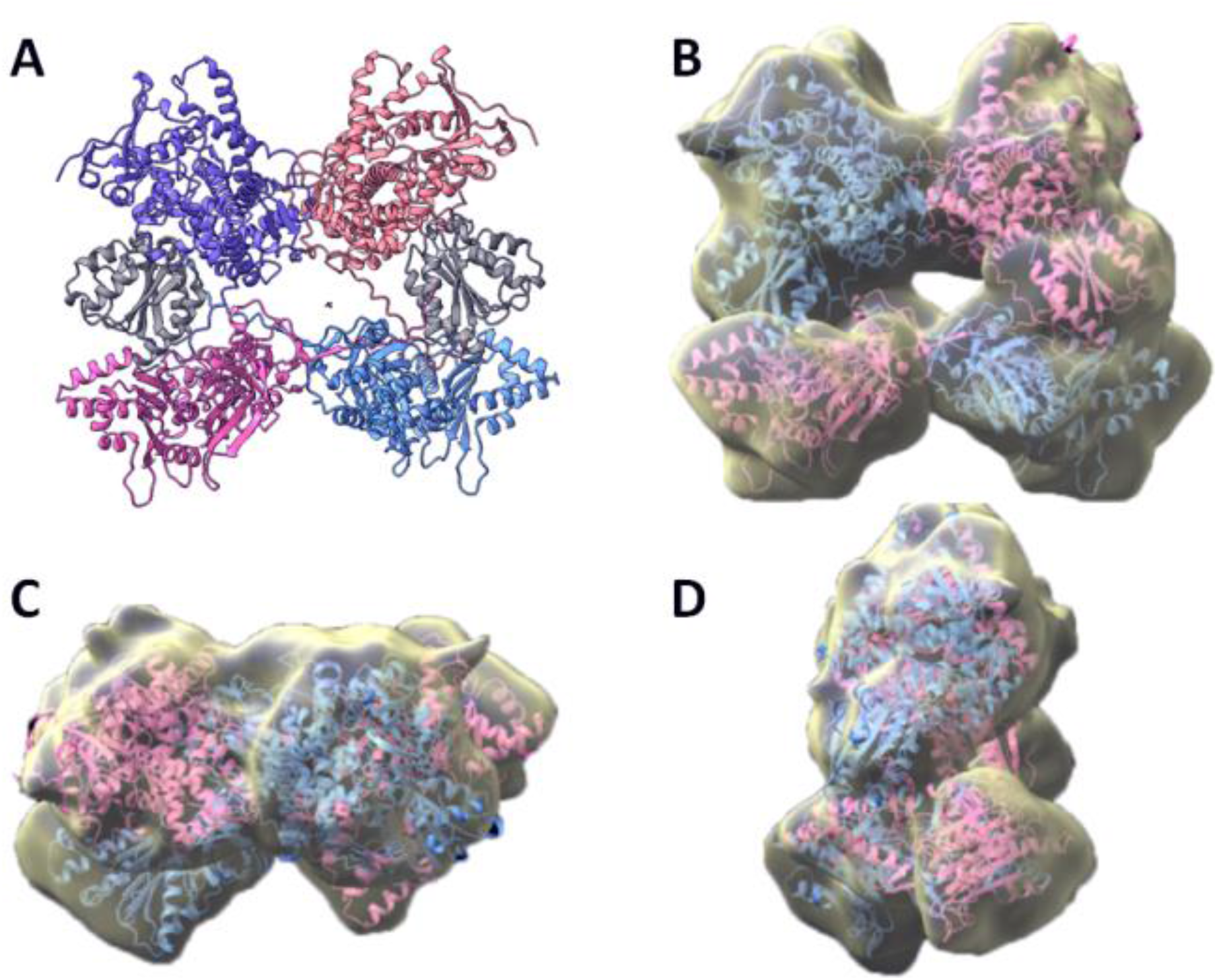
Comparisons of AF2A-predicted and cryo-EM experimental structures of full-length CYP102A1 dimer with the reductase domains in an open conformation. *Panel A*. The AF2A full-length model was generated by assembling partial structures for the predicted P450d-FMNd dimer and FADd dimer models as described in Methods. The two P450d are shown at the top, and the reductase dimer at the bottom. The salmon-colored P450d belongs to same chain as the pink-colored FADd, whereas the violet-colored P450d is on the same chain as the blue-colored FADd. The two FMN binding domains complexed to P450d are colored in grey. *Panels B, C, and D*. Model from panel A was embedded into the EMD-20785 density map and is viewed under different angles (face, bottom and side, respectively).

Comparison of models featuring reductase domains in the open and closed conformations (Supplementary Figure S6) illustrated that the two conformations are compatible with a similar positioning of the dimeric P450d and FADd parts within the same EM envelope. However, FMNd rotation required for catalysis can be sterically hindered within the limits of the two alternate models. Some structure relaxation, for example by a limited transient motion of the two contacting FADd in CYP102A1 dimer, would be greatly facilitated by the preliminary dissociation of the inter-chain FADd-FMNd complex and the rather low calculated stability of contacts at the FADd-FADd interface.

No direct contact exists between P450d and FADd in the closed conformation of the enzyme, in which position of the P450d dimer appeared only restricted by the possible extensions of P450d-FMNd linkers. An asymmetric hybrid conformation of the full-length CYP102A1 enzyme in which one of the two monomers would be in the open conformation and the other in the closed conformation was modeled (Supplementary Figure S7). Such rupture of symmetry would be consistent with the alternate cryo-EM images^22^ reported as partially opened structures. This geometry could favor a transient rotation of the P450d dimer around its symmetry axis facilitating FMNd flip-flop motion. This geometry also illustrates the possibility of complementary filling of the EM-20786 map by alternate FMNd conformations (right panel in supplementary figure S7). It is important to note that while the modeling by parts of the full-length CYP102A1 involved virtual polypeptide chain cleavages, reestablishing connectivity on presented fully assembled structures was always found geometrically feasible. However, following peptide chains linking in the assembled global structure some local remodeling of the linker geometries would be required for final improvement of the global model.

### 5. Determining mechanisms leading to dimer formation and design of synthetic dimeric monooxygenase

Mechanisms leading to the formation of the unusual CYP102A1 dimeric structure were questioned with the aim to design synthetic dimer structure of other P450 enzymes mimicking the high specific activity of CYP102A1. Dimerization of the P450 domain and dimerization of the reductase domain are two events that likely occurred independently during the evolution. To find traces of these events, the NCBI database was searched by PBLAST for sequence similarities with the reductase domain of CYP102A1 excluding its P450 domain from the query to avoid biasing the search. Surprisingly, the large number of sequence hits featuring more that 50% sequence identity with the CYP102A1 reductase domain were all bacterial natural fusions with a P450. The sequence collection was then filtered to remove duplicate or too similar sequences (> 98% amino acid identity). These filtered sequences were clustered to finally retain five typical sequences belonging to different microorganisms and exhibited variable similarity between them (from 54 to 95 % identity). The P450 domains found fused in the selected sequences exhibited from 64 to 98 % amino acid identity with the CYP102A1 P450 domain (Supplementary Table S2).

These five selected sequences were AF2A modeled by parts as described for full-length CYP102A1. All of them were predicted to share the same structural organization than CYP102A1, with an independent dimerization of the P450 domains and the possible formation of a reductase dimer stabilized by crossed FADd-FMNd complexes (data not shown). The calculated interfaces in these dimers and their binding free energies were estimated and found very similar between them and with those calculated for full-length CYP102A1. Interestingly, the P450d-P450d interfaces in the five selected sequences encompass 18 highly conserved residues that were individually predicted to be in contact in more than 50% of AF2A modeling attempts (Supplementary Figure S8). To explore whether this P450d dimerization constitutes a specificity of considered microorganisms, the three other P450 sequences present in the *Priestia megaterium* genome (WP_029321191.1, WP_013058569.1, and WP_053488633.1) were examined. In contrast to CYP102A1, these sequences correspond to bicomponent P450 systems. AF2A modeling indicated that none of them was predicted to dimerize. This suggested that dimerization of CYP102A1 and related enzymes constitute a specificity of bacterial P450-reductase fusions and not a particularity of P450 enzymes in the considered microorganisms.

A similar approach was designed to validate that the formation of a crossed P450-reductase dimer was not specific to the microorganism. *P. megaterium* genome was searched for the presence of other diflavin reductases similar to eukaryotic CPRs but not associated with a P450 domain. A unique sequence (WP_057244461.1) was identified that exhibited a higher similarity to assimilatory sulfite reductases than to classical CPRs. These two enzymes, sulfite reductase and cytochrome P450 reductase, belong to the same superfamily^31^, and Bernhardt *et al*. have shown that soluble sulfite reductase can also support the activity of a bacterial cytochrome P450, namely CYP106A1, and that of eukaryotic CYP21A2^32^. Possible dimerization of *P. megaterium* sequence WP_057244461.1 was not predicted during AF2A modeling. This raised the question of the molecular basis leading to the specific potency of bacterial CYP102A1 reductase domain to spontaneously dimerize. Considering the different types of interchain interactions deduced from AF2A modeling work presented above compared to the experimental structures of eukaryotic CPRs, three hypotheses were tested: i) presence of differential structural constraints on the orientation of the FMNd-FADd linker (including the hinge and the N-terminal long loop of the FAD domain), ii) some significant difference of length of the FMNd-FADd linkers between CYP102A1 and eukaryotic CPRs, iii) some differences in the geometrical complementarity at the FADd-FADd interface.

To test the first hypothesis, we considered the role of a small β-strand in the FMNd-FADd linker involving amino acid residues 644-647 of CYP102A1. This small β-strand constituted in CYP102A1 the entry point of the interdomain linker at the FADd-FADd interface. It stands immediately upstream of a characteristic turn present in all CPRs but absent in CYP102A1 reductase (See Fig. 8D). In eukaryotic CPRs, this loop goes back into the FADd (Loop 1 in Fig. 8D) whereas the presence of the β-strand in CYP102A1 (P1 in Fig. 8D) directs the chain toward the other monomer. It thus seems that the presence or the absence of this β-sheet alternatively redirects the chain into an intra-or an inter-molecular way. The native ^644^SLQF^647^ sequence of this β-strand was substituted *in silico* by the ^644^AGAG^647^ sequence in order to try to destabilize this fold and the corresponding reductase variant was AF2A-modeled. Two distinct types of folds, monomeric and dimeric, were randomly generated upon repeated AF2A modeling of the variant (Supplementary Figure S9). The modified 4-residue long sequence remained a β-strand in the dimeric structure while being converted into a non-structured region in the mutant monomeric structure. In the first type (Panel A), the FMNd is associated to FADd in an intramolecular complex, whereas in the second type (Panel B), the FMNd-FADd linker adopted a different orientation due to the 644-647 β-strand involved in a small β-sheet with another β-strand from FADd leading to formation of the native intermolecular complex. However, a demonstrative conclusion from this example must be mitigated by the fact that the efficiency of conversion from a dimeric to a monomeric structure appeared variable when testing alternate sequence substitutions. We thus concluded that destabilization of the β-strand at the FMNd-FADd linker at the FADd-FADd interface could favor the monomeric form of the reductase but did not constitute alone a sufficiently determining factor for the folding difference between CYP102A1 reductase domain and eukaryotic CPRs.

The second hypothesis, which establishes a link between the CYP102A1 structure and the length of the FMNd-FADd linker, was already previously proposed. The length of this linker region (hinge + N-terminal loop of the FADd) is systematically shorter in CYP102A1 and related enzymes compared to monomeric eukaryotic CPRs. The Figure 11 presents a multiple sequence alignment (panel A) of the FMNd-FADd linker regions between several eukaryotic CPRs and several bacterial CYP102A1-related proteins. As illustrated, the linker in bacterial reductase domains, that systematically form a dimer, is systematically shorter by about 20 amino acids than the corresponding region in eukaryotic CPRs, which are all systematically monomeric. From this observation, we devised a first CYP102A1 mutated sequence in which the FMNd-FADd linker was extended by introducing 6 amino acid residues GSGGSG, forming a highly flexible region, in front of the N-terminal part of the linker, transforming the ^648^VDSAADM^654^ wild-type sequence in the longer GSGGSGVDSAADM mutated sequence. Reciprocally, a second mutated sequence was derived from the human CPR sequence in which the ^254^DAAKVYMG^261^ sequence spanning the aligned region specific of monomeric CPRs was deleted, transforming the wild-type ^250^HTDIDAAKVYMGEMGRLK^267^ in the shorter mutated sequence HTDIEMGRLK. As shown in the panel B, the positions of the exchanged sequences were based on structure-directed sequence alignments to compensate for the limited sequence similarity in these regions.

Sequence pairs of the two resulting mutants were AF2A modeled. Figure 12 shows that the 8-residue deletion in human CPR virtual mutant did not significantly modify the predicted CPR folding except for some structuration of the shortened loop with a supplemental short α-helix (panels A and B). No dimer formation was predicted upon repeating modeling (up to 10 runs). In sharp contrast, a major conformational change was observed when the FMNd-FADd linker of the bacterial enzyme was extended by 6 residues. The insertion converted CYP102A1 reductase domain from a dimeric form to a strictly monomeric form similar to monomeric eukaryotic CPRs (panels C and D). This result based on *in silico* mutations clearly suggests that the shortened FMNd-FADd linker systematically found in dimeric bacterial enzymes is critical for dimer formation. Considering the absence of reciprocity upon linker extension in the monomeric CPR, this suggested that the role of the shortened linker was to destabilize the formation of intrachain complex thus redirecting the folding to a crossed dimeric structure as observed in all bacterial P450-reductase fusion enzymes. In contrast, observation that shortening of the CPR linker was inefficient to promote dimerization suggested the requirement of some supplementary factors.

**Figure 11.**
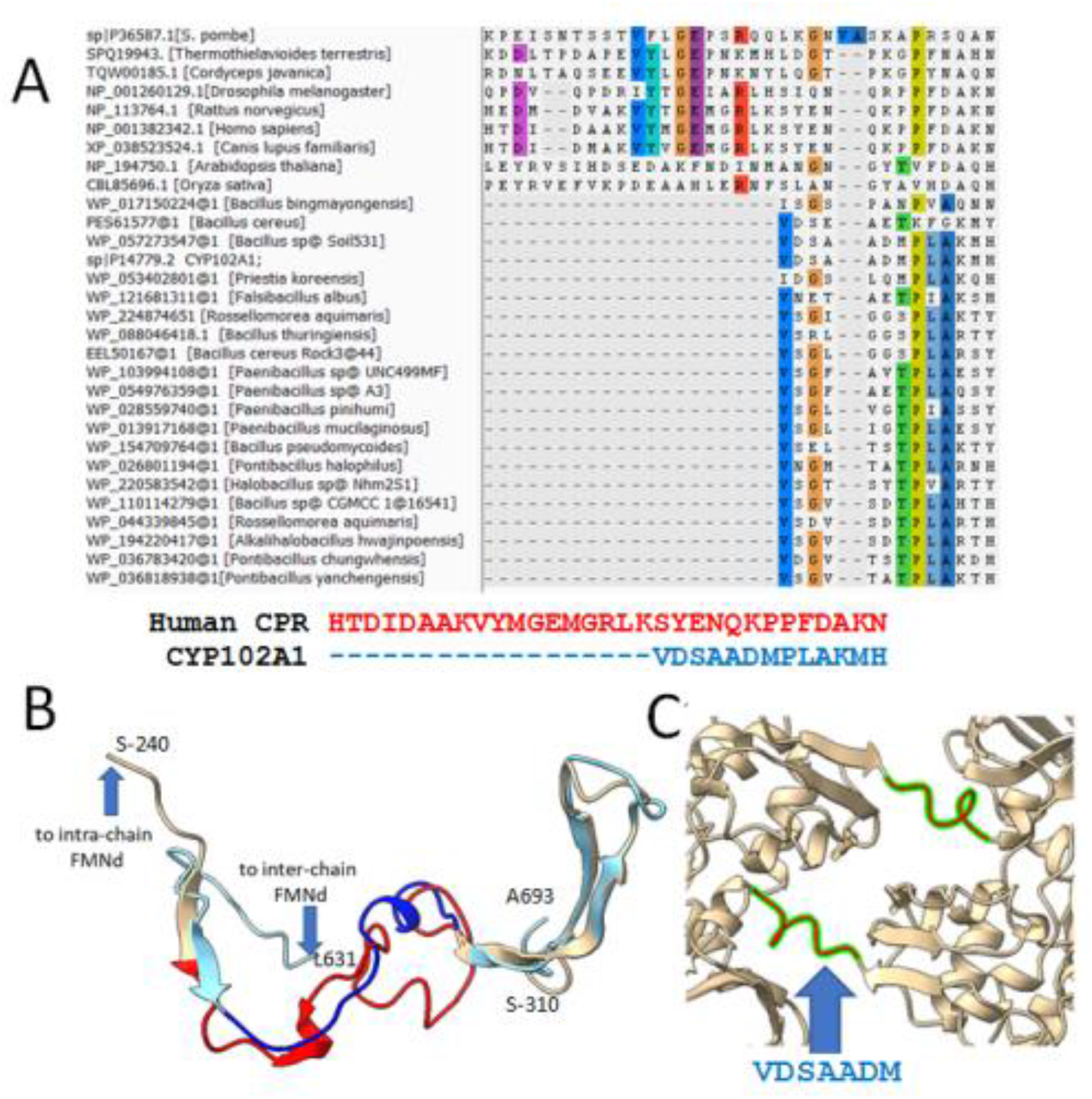
Alignment of the FMNd-FADd CYP102A1 interdomain linker region and of the corresponding eukaryote CPR sequences and AF2A modeling of *in silico* mutated sequences. *Panel A*. Due to limited sequence similarities in this region between bacterial and eukaryotic sequences, alignment limits were based on common motifs in the respective structural models and encompasses the two common β-strands present in the structures of human CPR and *P. megaterium* CYP102A1. *Panel B*. Structural overlay of the corresponding regions. Crystal structure of human CPR (PDB 3qe2) is colored in wheat. It extends on this representation from Ser240 to Ser310, with the critical loop ^250^HTDIDAAKVYMGEMGRLKSYENQKPPFDAKN^280^ colored in red. The corresponding segment in the CYP102A1 monomer is colored in pale blue with its shorter loop ^648^VDSAADMPLAKMH^660^ colored in dark blue. Monomeric structures were superposed by RMSD minimization restricted to the FAD binding domain. Common β-sheet motifs used for alignment are visible on the right and left part of the mutated longer (in red) and shorter (in deep blue) sequences. *Panel C*. Structure of the ^648^VDSAADM^654^ bacterial peptide (colored green-red) in the AF2A model of CYP102A1 reductase dimer.

**Figure 12.**
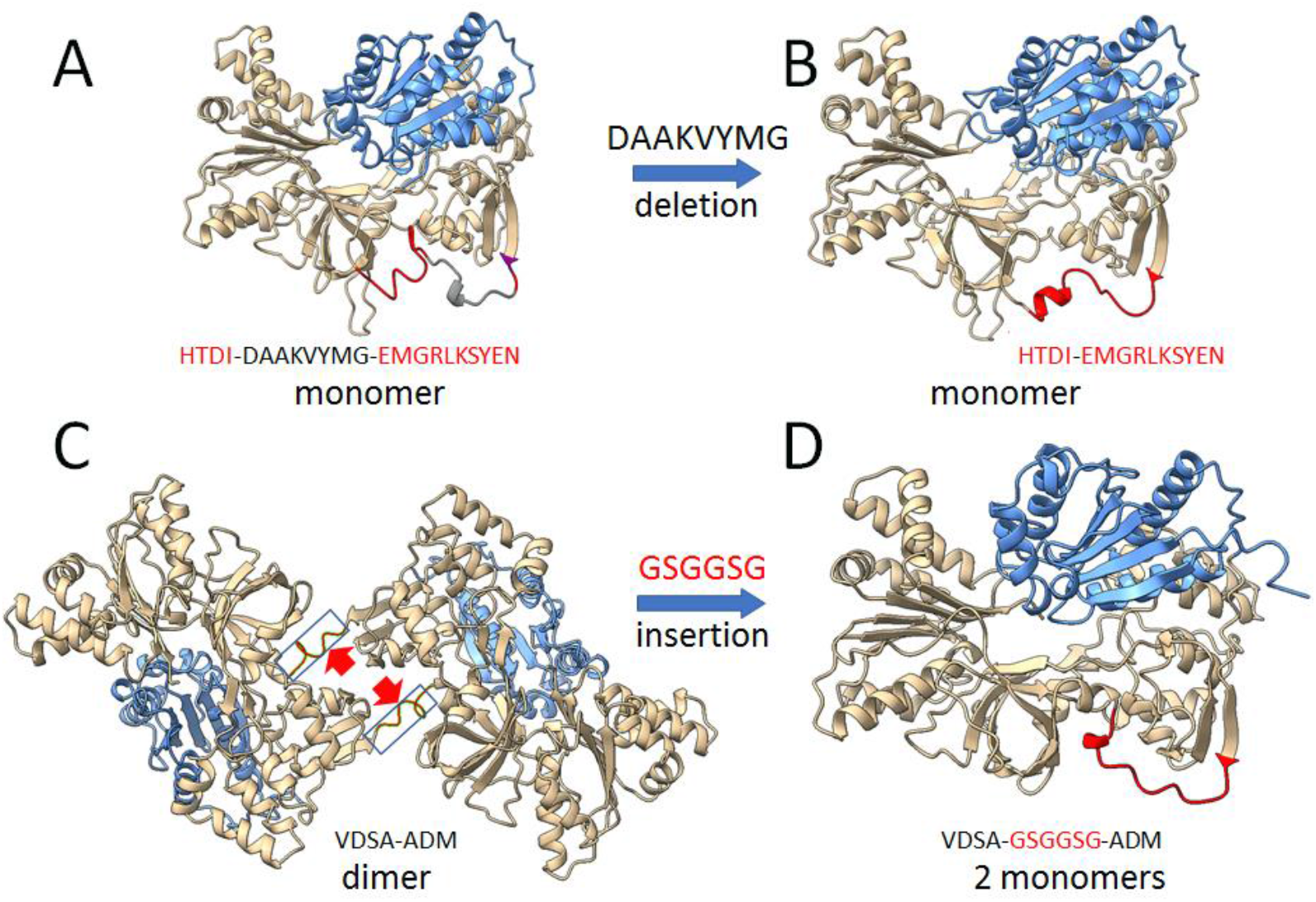
Reverse-engineering of human CPR and CYP102A1 reductase domains. *Panels A and B*. AF2A prediction of the structural consequence of the deletion in human CPR of the ^254^DAAKVYMG^261^ sequence in the loop connecting FMNd to FADd (this loop is directly C-terminal to the hinge segment). *Panel A*. Structure of the wild-type, native human CPR. Red and grey colors indicate the correspondence between targeted sequence and structure. *Panel B*. Structure of the DAAKVYMG deleted mutant of human CPR. Targeted region is visualized in red. The insertion did not cause major structural change, the model remaining monomeric. *Panels C and D*. Predicted structural consequence of the insertion in CYP102A1 of the GSGGSG sequence in the middle of the ^648^VDSAADM^654^ sequence. *Panel C*. Structure of the wild type reductase dimer. The framed area indicates the VDSAADM segment and the red arrows indicate the point of insertion of the GSGGSG sequence. *Panel D*. AF2A-predicted structure of the mutant with the extended FMND-FADd linker colored in red. Conversion of the dimeric structure into a monomeric structure occurred following such insertion.

The third hypothesis proposed concerns the role on dimerization of the shape and charge complementarity at the FADd-FADd interface. To evaluate this role, a chimeric human-bacterial ‘Janus’ sequence was designed in which the sequences of the FMN domain and the FMNd-FADd interface were those of human CPR, while the sequence of the FADd-FADd interface was that of CYP102A1 reductase domain. Such a ‘Janus’ design raised the difficulty of maintaining the structural integrity at the junctions between the CPR and bacterial sequences in the final chimeric amino acid sequence. The proof of concept was designed as to preferentially target all these junctions into the more conserved structural regions, as is observed with chimeric enzymes produced randomly in yeast^33^. However, a more careful design and local sequence optimizations should likely be required to facilitate transposition of this *in silico* approach at experimental level. Feasibility was evaluated for a chimera where five sequence segments of human CPR were substituted by the corresponding sequences of the CYP102A1 reductase domain (Supplementary Table S3). This design was based on the previously defined requirements for the structure of a Janus enzyme. The resulting synthetic chimeric sequence was assayed for dimer formation with AF2A. While the majority of resulting models were still folded as pairs of independent monomers indicating the absence of dimer formation, in 2 cases out of 10 (20%) AF2A successfully generated a well-defined dimer folded in the crossed configuration typical of CYP102A1 (Figure 13). Such score was significantly lower than the success rate (80% of modelling attempts) for dimer prediction with the native CYP102A1 reductase sequence, but was in the same range than the 20% value obtained for the dimeric geometry of the PES61577 sequence from *Bacillus cereus* (See supplementary Table S2), a 1,049-residue long protein that shared 68% amino acid identity with CYP102A1. The predicted dimeric structure of the chimeric ‘Janus’ sequence was tightly mimicking the structure of CYP102A1 reductase domain dimer at the level of the interface between the two FAD domains. However, outside of this region, some differences were observed in loops and details of the structure of the FMNd-FADd complex both when comparing the AF2A models (Supplementary Figure S10) and comparing the hybrid structure model with crystal structures (Supplementary Figure S11). In the model of this synthetic sequence, the structure of FMNd was strictly conserved compared to the human CPR, as well as the structure of the FMNd-FADd interface but, as expected, significantly differed from the structures in the bacterial counterpart.

**Figure 13.**
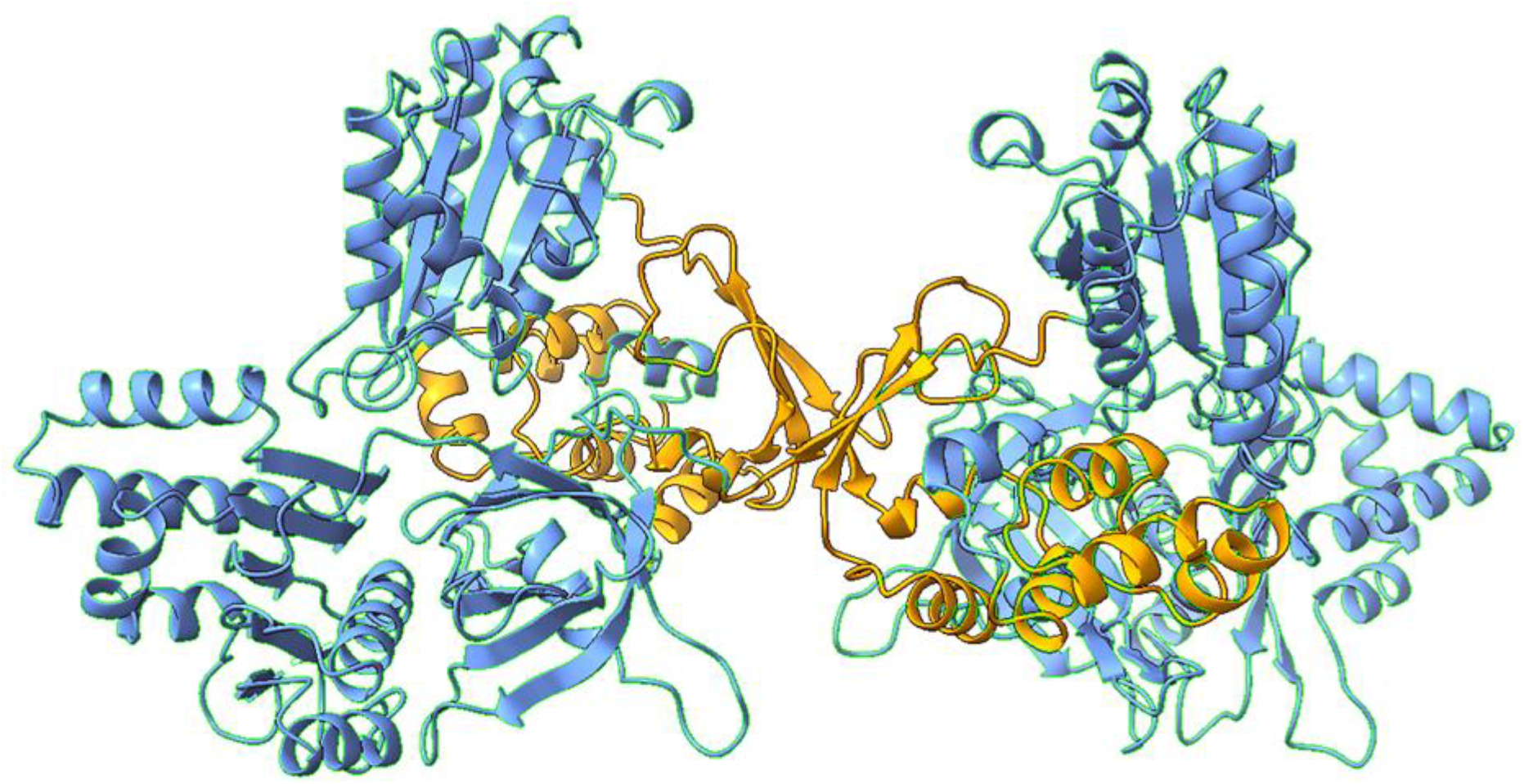
AF2A predicted dimeric structure for a SYN01 mosaic human-bacterial reductase domain. The two FMNd and the FMNd-FADd interfaces are constituted of human sequences and are colored in blue, the FADd-FADd interface is constituted of bacterial sequences and is colored in yellow.

As summarized in supplementary Table S4, the structural factors associated with the three hypotheses presented above played complementary roles in dimer formation. This engineering route thus appeared extremely interesting for the design of a synthetic system with dimeric eukaryotic CPRs in which FMNd and its interface to FADd would be made optimal to efficiently transfer electrons to any fused eukaryotic P450 in various biotechnological applications. Optimization of chimeric reductase construct based on the sequence of a eukaryotic CPR, followed by its fusion to a P450 of the same species, targeting the exceptionally high turnover permitted by the CYP102A1 dimeric crossed structure, would constitute a promising starting point to optimize monooxygenase activity of any P450-CPR artificial fusions.

We developed an *in silico* proof-of-concept that a mosaic reductase element, combining sequences of eukaryotic CPR and CYP102A1 reductase domain, able to self-dime--rize can be designed. Such reductase dimer could be fused to potentially any P450 of interest. Reproducing the geometrical complementarity of bacterial P450d-P450d interface with heterologous P450s without affecting their functions would be an extremely challenging task. In contrast, experimental optimizations, for example by directed evolution of the *in silico* designed animal or plant synthetic dimeric reductase domain appeared a much easier task. Consequently, transferring the high turnover properties of native CYP102A1 into synthetic dimeric P450-CPR fusions would be simplified in the case where affinity between P450d domains would be only accessory for dimerization and for the formation of the crossed structure at reductase domains interface.

### 6. Modeled CYP102A1 crossed structures resolve contrasting biochemical data

Considering AF2A and previously reported models, CYP102A1 dimer can involve four possible alternate topologies resulting from the presence or the absence of chain crossing at the levels of P450d-FMNd and the FMNd-FADd connections. Neeli *et al*.^16^, then Kitazumi *et al*.^17^ two years later addressed this question by biochemical analyses of CYP102A1 site-directed mutants. They designed four mutations, F87Y and A264H, G570D, and W1046A that specifically inactivate the function of the P450 domain (for the two firsts), the FMN domain, and the FAD domain, respectively. A mixture of homodimers and heterodimers were reconstructed by associating polypeptide chains carrying individual or pair of the above mutations and the effects of these mutations on NADPH-supported fatty acid hydroxylase activity and on NADPH-cytochrome *c* reductase activity were analyzed. Based on our AF2A models, we calculated the CYP102A1 residual activity for the four possible combinations of mutated polypeptide chains and compared these results to the experimental values previously reported (Figure 14). Calculations assumed that the monooxygenase activity required that functional FADd, FMNd and P450d be simultaneously present on the same side of a symmetrical dimer structure thus permitting electron transfers. Full dimer dissociation into monomers allowing exclusive intrachain electron transfers was excluded, since they were not observed during cryo-EM image acquisition^22^, and since monomeric enzyme were found to be inactive^16^. The two-electron transfer chains present in each monomer of the native dimeric enzyme were also considered to equally contribute to activity and to be able to work independently. We previously illustrated on supplementary Figure S5 that, due to the symmetry, both crossed and non-crossed P450d-FMNd connectivity are geometrically possible. An additional model for mutant was thus considered assuming that a CYP102A1 dimer can randomly harbors the two FMNd-P450d configurations with arbitrarily similar probabilities. Residual monooxygenase activities expected for the four combinations of mutated chains and the hybrid configuration described on Figure 14 were calculated. To do that, we assumed that activity of the dimer was the sum of the contributions of the two possible electron transfer chains and that mutation of a domain interrupted its corresponding electron transfer without affecting global CYP102A1 domain organization or dynamics. Comparison of predictions with reported experimental values clearly excluded the association of a non-crossed topology of FMNd-FADd (indicated as NCB) with the non-crossed topology of P450d-FMNd (indicated as NCP). This topology would be completely inactive in three mutant combinations (A-B, B-C, C-D) where experimental activity was indeed observed.

**Figure 14.**
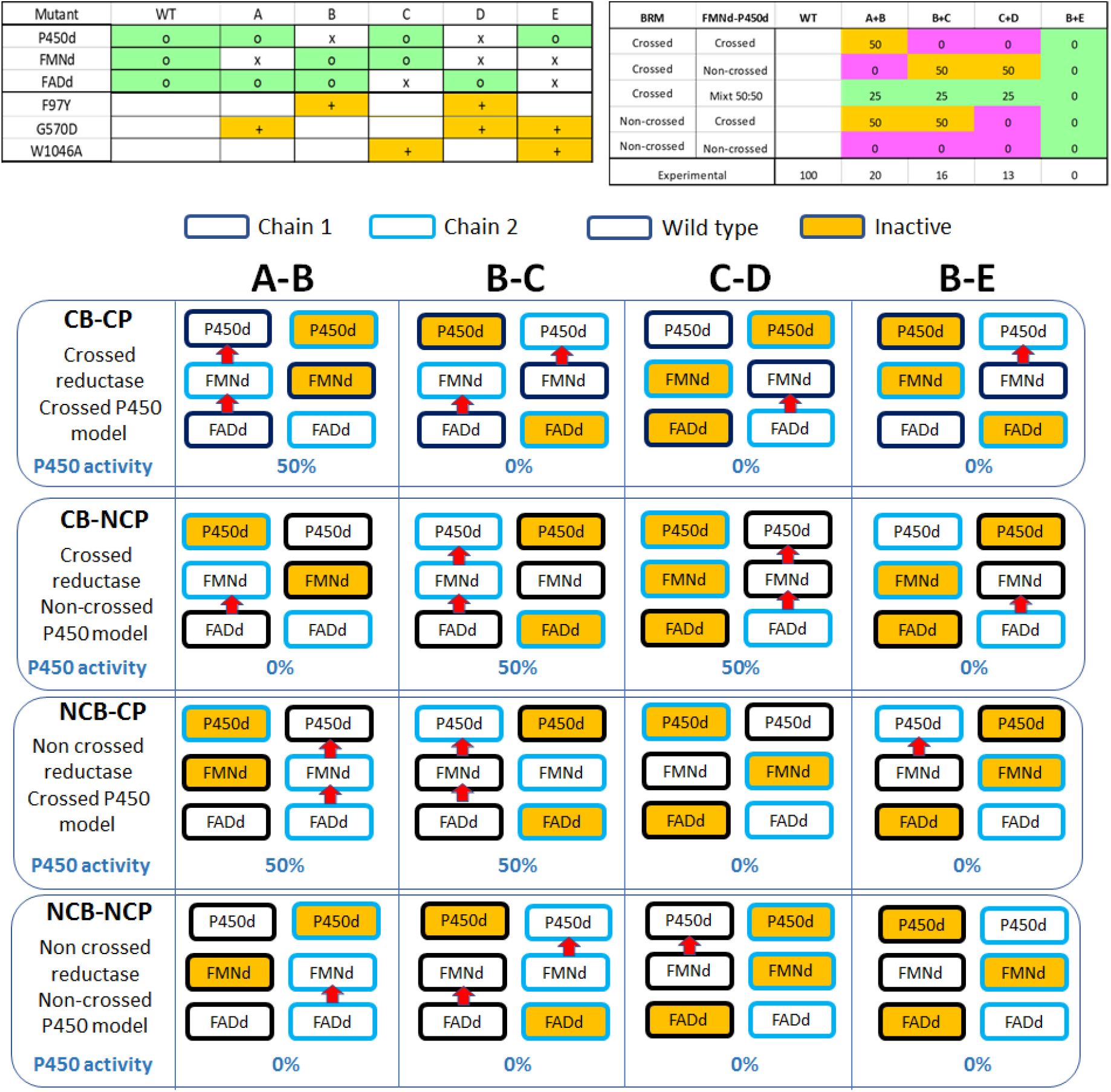
Comparison of CYP102A1 predicted fatty acid hydroxylase activities assuming different topologies with published experimental data. Five different sets of literature described mutations (A-E) inactivating either the P450d, FMNd and FADd functions or pair of these functions are illustrated on top panel. Yellow boxes indicate mutation(s) and green boxes the corresponding functional consequences in each case. CYP102A1 residual activities reported for reconstructed heterodimers associating A-B, B-C, C-D and B-E mutant chains have been reported^16,17^ and are given (in per cent of the wild-type activity) in the ‘Experimental’ row on the bottom of the top right panel. Residual activities corresponding to these combinations of mutations have been predicted considering the four possible topologies for CYP102A1 dimer involving crossed (CB) or non-crossed (NCB) FMNd-FADd dimer and crossed (CP) or non-crossed (NCP) FMNd-P450d dimers. These four possible topologies in association with the four types of described associations of mutations are illustrated on the bottom panel. In this panel, yellow-colored boxes correspond to inactive domains. The border color of each box (black or blue) labels domains belonging to the same polypeptide chain considering each topology. Residual activity in each case was calculated assuming that a full electron transfer chain (indicated by red arrows) from FAD to P450 occurring on a same structural side of a dimer was required for half of the full dimer functionality and that alternate (left or right) ways are equivalent and functionally additive. Predicted results are indicated in each case (topology / mutant combination) in the row ‘P450 activity’ in percent of the wild type value. Theoretical values are summarized into the top-right panel to be compared to experimental data.

The situation in which both P450d-FMNd and FMNd-FADd topologies are simultaneously crossed (respectively indicated as CP and CB) was inconsistent with experimental values in two cases. Mutated chain combinations B-C and C-D would be inactive in contrast to experimental data. The two other situations (CB-NCP and NCB-CP) also appeared inconsistent with the experimental data. The NCB-CP topology retained in the cryo-EM study^22^ was inconsistent in one case (the C-D mutant) and would lead us to predict a residual activity must larger than the experimental value in two others cases (A-B and B-C mutants). The last case (topology CB-NCP) was similarly incompatible in one case (A-B mutant) and hardly compatible in two cases (B-C and C-D mutants) leading to the absence of any simple solution. However, AF2A modeling suggested that CP and NCP topologies can coexist. The hybrid topology (CB-NCP/CP), in which the reductase domains are in a crossed configuration with the P450d-FMND complex being partly in a crossed configuration and partly in a non-crossed configuration was found to be consistent with experimental values. For the four considered combinations of mutated chains, a residual activity of heterodimers fairly close to the experimental values was predicted (theoretical 25% compared to experimental values of 20%, 16% and 13%). Parasitic formation of small amounts of homodimers between mutated chains giving non-functional enzymes lacking the possibility of the *trans*-complementation can be statistically expected during experimental hybrid dimer reconstitution. This would yield some excess of fully inactive dimers that could explained the fact that observed activities remained lower than theoretical values deduced from AF2A models. We thus concluded that the sole topology consistent with the reported experimental effects of mutant chain combinations was the ones involving a crossed reductase dimer and a mix of crossed and non-crossed FMNd-P450d configurations. This conclusion is consistent with the requirement of an obligatory intermolecular FAD to FMN electron transfer as reported by Neeli *et al*.^16^.

In the crossed P450d-FMNd complex, the electron transfer from FMN to heme iron is intermolecular, as reported by Kitazume *et al*.^17^, while in the opposite case, this electron transfer would be intramolecular. In the cryo-EM study, the shortening of the length of the FMNd-P450d linker demonstrated to improve image quality, likely by reducing the interdomain dynamics, which in turn progressively inactivated the enzyme^22^. This observation can be interpreted with our AF2A model considering that linker length reduction would impair the freedom of rotation of FMNd that is permitted under the local constraints of two peptide segments, the linker with FADd that stabilizes an intermolecular arrangement for the FMNd-FADd complex, and the linker with P450d that stabilizes almost equally an intramolecular or an intermolecular P450d-FMNDd complex.

The autonomous dimerization of isolated P450d in AF2A predictions was also a source of contradiction with some previous experimental report. Size-exclusion gel chromatography clearly showed that purified isolated P450 domains of CYP102A1 were eluting as monomers^16^. No peak corresponding to the exclusion volume of a P450d dimer was observed. In contrast, the cryo-EM deduced models supported P450d dimer formation. Calculated binding free energy of about -9 kcal/mol deduced for dimer formation in our AF2A models corresponds to an expected binding affinity in the micromolar range that could be insufficient to promote dimer formation in gel filtration studies. However, when this P450d-P450d association is put in combination with the contribution of the binding energy resulting of the crossed reductase domains, this amounts to some -30 kcal/mol of binding free energy. The dissociation constant in the nM range experimentally observed for the dimerization^16^ can be easily explained by this finding from AF2A models.

## Concluding remarks

The present report illustrates that, in addition to accurate structural prediction, a poorly documented property of AF2A is its ability to provide alternate complex geometries. Considering that AF2A does not primarily involve energetic calculations, the frequency of observation of alternate geometries is difficult to link with alternate structure stabilities. However, we develop in this report an approach based on the competition for the formation of alternative and exclusive complexes that brings significant clues on the possibility of alternative complexes formation including those in response to minor sequence changes. The theoretical basis of such response is to our knowledge unclear in the AF2A algorithm and might represent the selection of preferential folding path resulting from geometric criteria. Recognized successes of the algorithm raised the question of the respective role of thermodynamic criteria and of geometric criteria that can in turn dynamically drive protein folding without requiring extensive sequence space exploration.

These properties were validated using CYP102A1 as a test case, taking profit of the large amount of biophysical and biochemical data available for this P450 enzyme. Confrontation of AF2A predictions with cryo-EM and crystallographic data was particularly instructive, notably by allowing us to correct and extend previous interpretations and particularly to propose a unified mechanism reconciliating existing experimental data. More generally, the stepwise model reconstruction developed by exploiting the evidenced AF2A properties can constitute a generally applicable *in silico* approach to the determination of alternate structure and dynamics of major biological complexes.

## Methods

### Softwares

AF2A modeling was performed using the Aphafold2_advanced Python notebook^34^ that was run on Google Collaboratory cloud computing facilities using GPU and its large (48 Go) dedicated memory. Google Colab Notebooks were run using default parameters except for the selection of the pTMscore ranking procedure and the num_models parameter that was adjusted between 5 and 20 depending on cases. Resulting models were visualized and analyzed using ChimeraX and Pymol softwares. Matchmaker and ISOLDE plugins in ChimeraX were used respectively for RMSD minimization and for model docking into cryo-EM density maps. AliView algorithm was used for sequence manipulations and visualization and the MIA-Muscle plugin for sequence alignments. TCoffee from Expresso web server was used for structure-based sequence alignments and the Prodigy (PROtein binDIng enerGY prediction) webserver was used to calculate the binding free energies (ΔG) of interacting domains. ConsRank and CoCoMaps web applications were used respectively to rank models of complexes and to generate interaction maps of contacting residues.

### Selection of models among alternate AF2A predictions

Each predicted structure (from 5 to 20) was examined. Structures in which domains were separated or overlapping were considered to sign the absence of any predicted complex. Overlaps, when present, did not affect individual domain folds and were never observed with domains belonging to the same polypeptide chain. When formation of a defined complex was predicted, the use of the AF2A recommended pTMscore was not systematically consistent with calculated binding affinities using the Prodigy software. In that case, the retained structure was based only on this last energetic criterion (i.e. the best affinity). When alternate geometries were predicted, the optimal fold for each was individually selected as previously. Formation of a dimeric structure was considered significant when predicted in at least 20% of the AF2A runs.

### Docking of AF2A models in the cryo-EM density maps

Electron density maps were filtered to a density level mostly eliminating solvent noise when fitting of the global shape of models was required. However, when structure details were targeted, filtering was increased to the higher-level improving resolution without visually impacting polypeptide chain continuity. Such levels can be different depending on the map considered and its resolution, and on the region targeted due to inhomogeneous levels of experimental map resolutions. AF2A predicted structure parts were manipulated as rigid bodies and not subjected to adjustment during docking into cryo-EM maps. For reconstruction of the full dimer structures, the two considered partial structures (either P450d-FMNd and FADd dimers or P450d and FMNd-FADd dimers) were first moved in a way that the symmetry axis of the P450d dimer became aligned with the symmetry axis of the FADd dimer. Then, distance between the two sub-models along this axis was reduced to the minimal value not creating clashes. The optimal rotation angle along this axis for the two sub-models cannot be determined by the *ab initio* modeling due to the absence of direct contact between the FADd and P450d and the expected alternate positioning of the FMNd to form complexes with the P450d and FADd. ChimeraX tools and its ISOLDE plugin were used to optimally define this angle using the experimental cryo-EM maps as a reference. During this process, structure fitting of the FMN domain into the cryo-EM density map was not considered to be a relevant criterion. Indeed, alternate complex geometries could well explain the significantly enlarged experimental cryo-EM density map at the FMNd location compared to the FMNd occupancy. Consequently, positioning of this domain was determined based on the P450d density fitting for the conformation involving the P450d-FMNd complex and based on FADd density fitting for the conformation involving the FADd-FMNd complex. This problem did not affect the fit for the isolated FMNd-FADd crossed complex. Due to the procedure of this reconstruction by parts, the continuity of the polypeptide chain must be reestablished after assembling the different sub-models in the cryo-EM density map. Structural feasibility was always checked and found possible but no energy minimization of the presented models was made in this work. All the complementary methods used to build each figure were detailed in respective figure legends.

## Abbreviations

AF2A: AlphaFold2_advanced
cryo-EM: cryo-electron microscopy
FADd: CYP102A1 FAD-binding domain
FMNd: CYP1021A1 FMN-binding domain
P450: cytochrome P450;
P450d: CYP102A1 heme-binding domain
SiR: sulfite reductase.

## Data availability

Atomic coordinates of the different AF2A-predicted CYP102A1 models are available by emailing the authors.

## Acknowledgements

We thank L. Garcia-Alles at TBI for valuable discussion and critical reading of the manuscript.

