## Supplemental Material for "Confrontation of AlphaFold2 models with cryo-EM and crystal structures enlightens alternate geometries of the CYP102A1 multidomain protein"

**Supplementary Figure S1.** Alternate AlphaFold2_advanced outputs when modeling complexes. *Panel A*. Proteins not forming a complex are independently modeled and their resulting structures randomly superimposed or juxtaposed. *Panel B*. Tightly related models are generated when two proteins can form a unique possible complex. *Panel C*. Structures for alternate possible complexes are generated when two proteins are competing for a common binding site on a third partner. The protein not forming a complex is then randomly placed in the model. *Panel D*. Structure for a ternary complex is generated when three partners exhibit independent interaction sites.

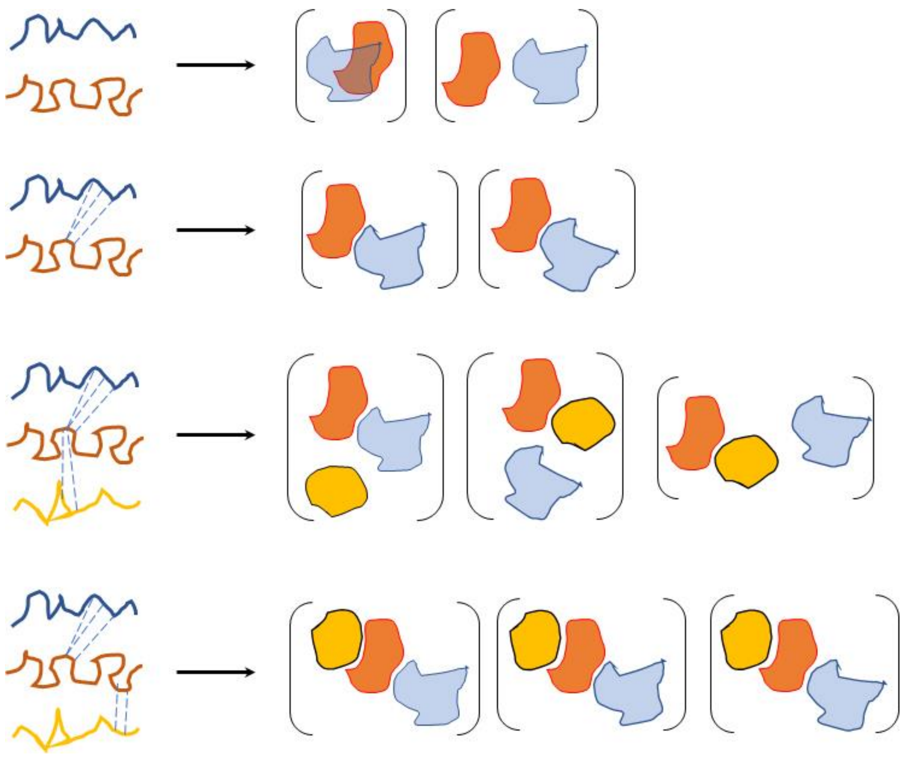

**Supplementary Figure S2.** AF2A predicted structures for the monomeric full length CYP102A1. *Panel A*. Model from the AlphaFold Protein Structure Database (ADB entry P14779). *Panels B, C, D*. Alternate AF2A predictions from repeated runs. Relative orientations of the four models were adjusted for comparison purpose using orientation of the P450 domain (CYP102A1 residues 1-476) of the ADB P14779 structure as a reference. P450d is colored in red, FMNd in yellow, and FADd in blue.

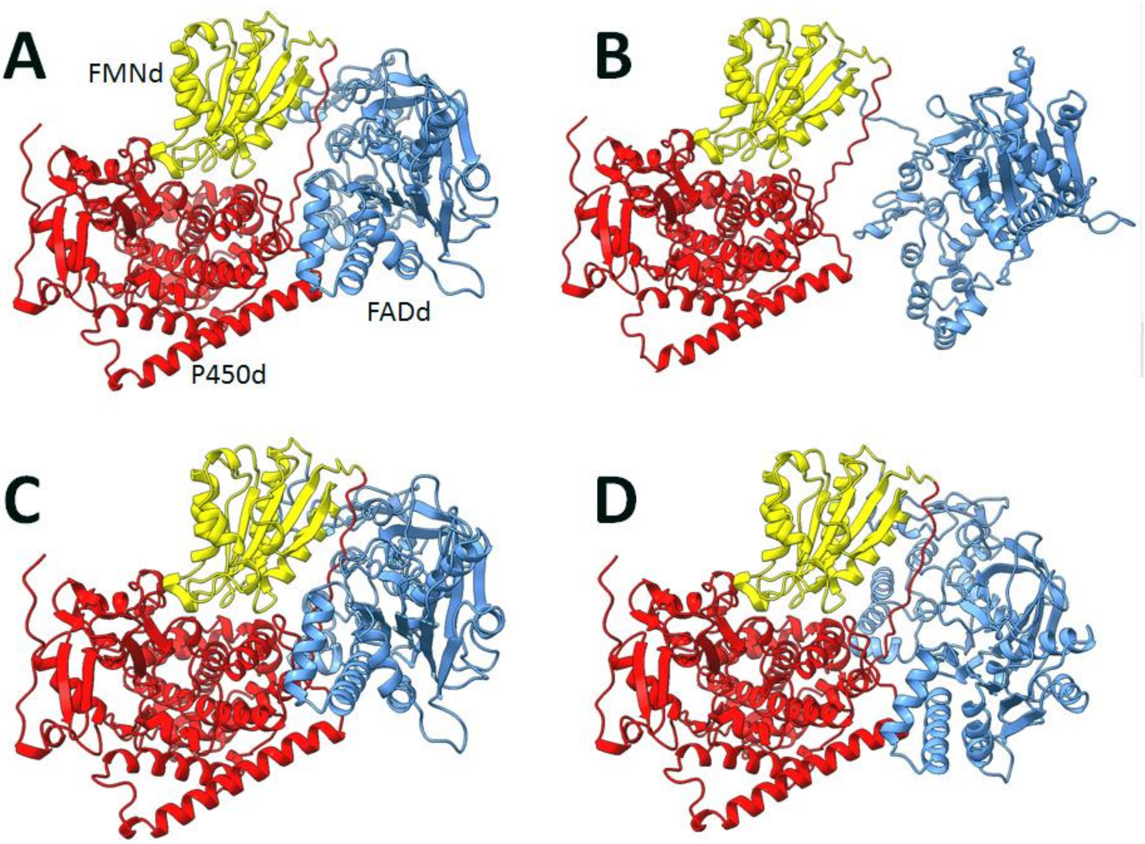

**Supplementary Figure S3.** Predicted cofactor geometries based on AF2A-based open and closed conformations of the CYP102A1 reductase domains. Cofactors were replaced into AF2A structure by cofactor atom coordinates transfer following RMSD minimization of AF2A structures with reference crystal structures of individual domains (P450d, FMNd, FADd). *Panel A*. FMNd-FADd interface in the crossed FMNd-FADd dimer in the closed conformation. *Panel B*. P450d-FMNd interface in the crossed FMNd-P450d dimer with the reductase domain in the open conformation. The cysteine ligand of the heme iron is also visualized in stick model. The heme and flavin cofactors are represented in stick models and colored yellow for the flavins and red for the heme.

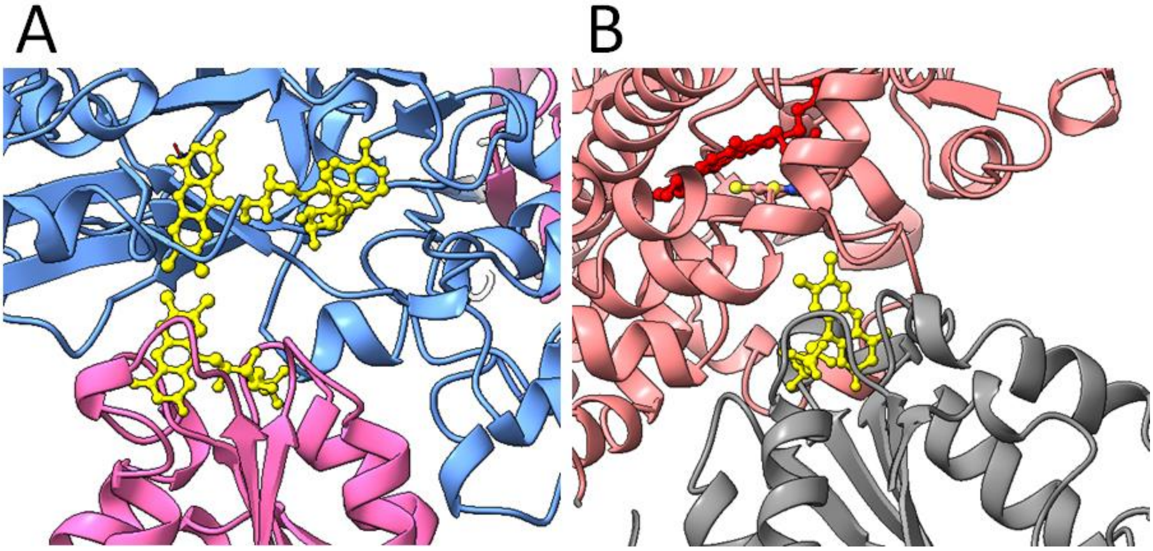

**Supplementary Figure S4.** Structural overlap between the PDB 1bvy crystal structure of CYP102A1 that includes two P450d and one FMNd per asymmetric unit (colored wheat), and the AF2A-predicted structure of the P450d dimer colored in pale blue. The FMNd domain appears complexed with one of the P450d in the crystal.

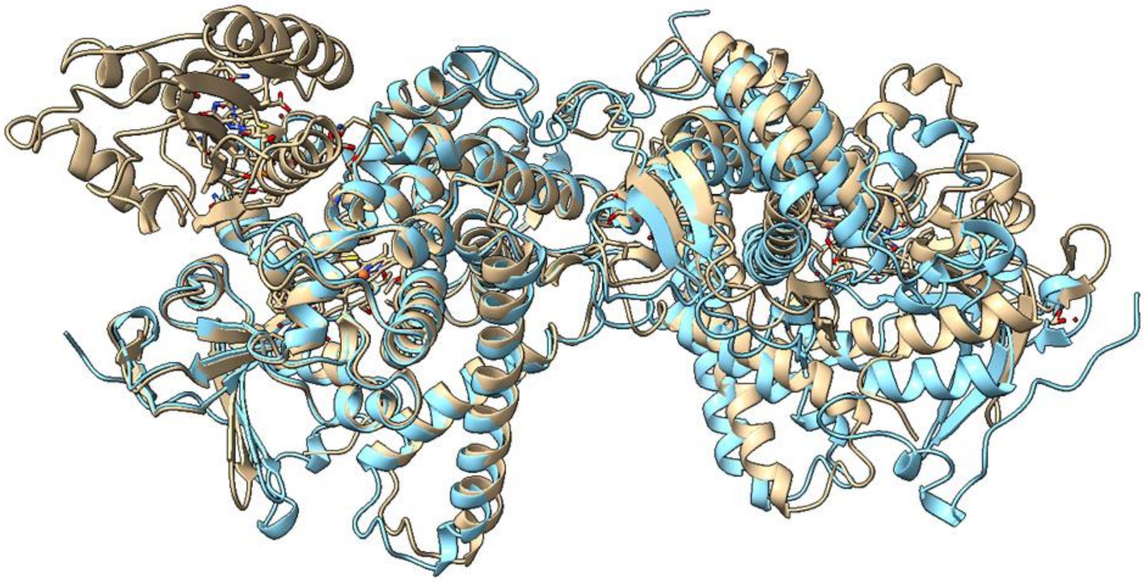

**Supplementary Figure S5.** AF2A predicted structures for the *cis* and *trans* conformations of the dimer formed from a pair of P450d-FMNd chains. In both cases, P450d and FMNd adopted similar complex geometries compatible with electron transfers. The *cis*-conformation, in which the complexed P450d and FMNd belong to the same chain, is illustrated by chains in blue and green colors. The *trans*-conformation, in which the complexed P450d and FMNd belong to the different chains, is illustrated by chains in the red and wheat colors.

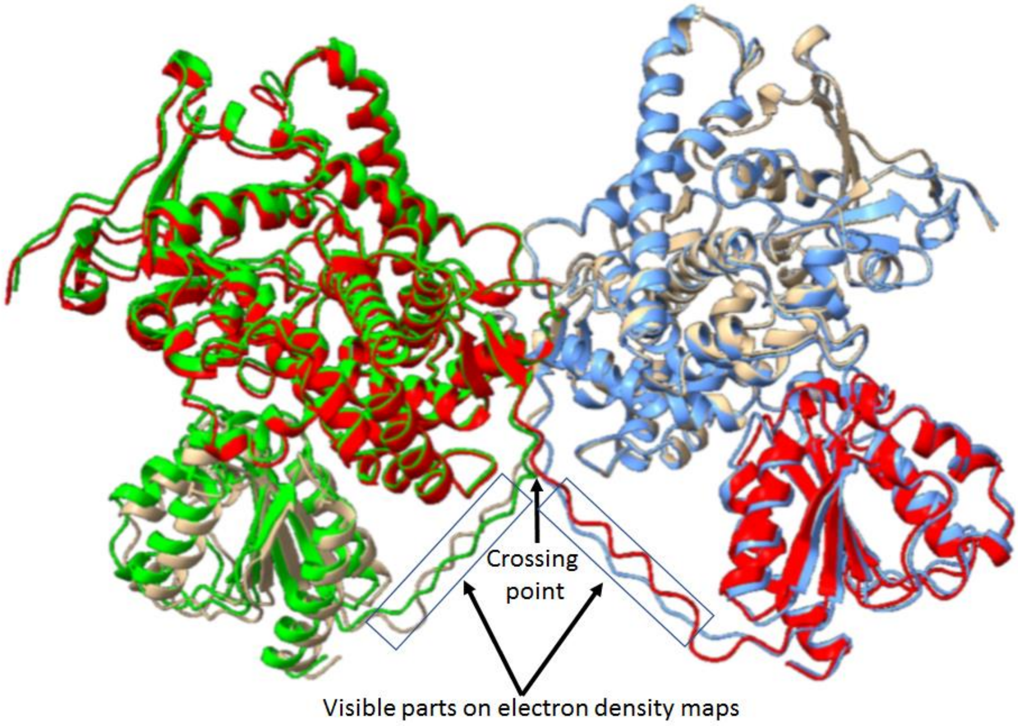

**Supplementary Figure S6.** Comparison of AF2A-predicted structure of full length CYP102A1 with the reductase domains in closed conformations (at left) or in open conformations (at right). The P450d are at top and the FADd at bottom and the FMNd in between. For the sake of clarity, the FMNd of each interacting monomers are white colored to help visualization of their different orientations in both configurations. P450d and FADd are colored blue for one monomer in the dimer, and pink for the other monomer. In the two configurations, FMNd and FADd are cross complexed.

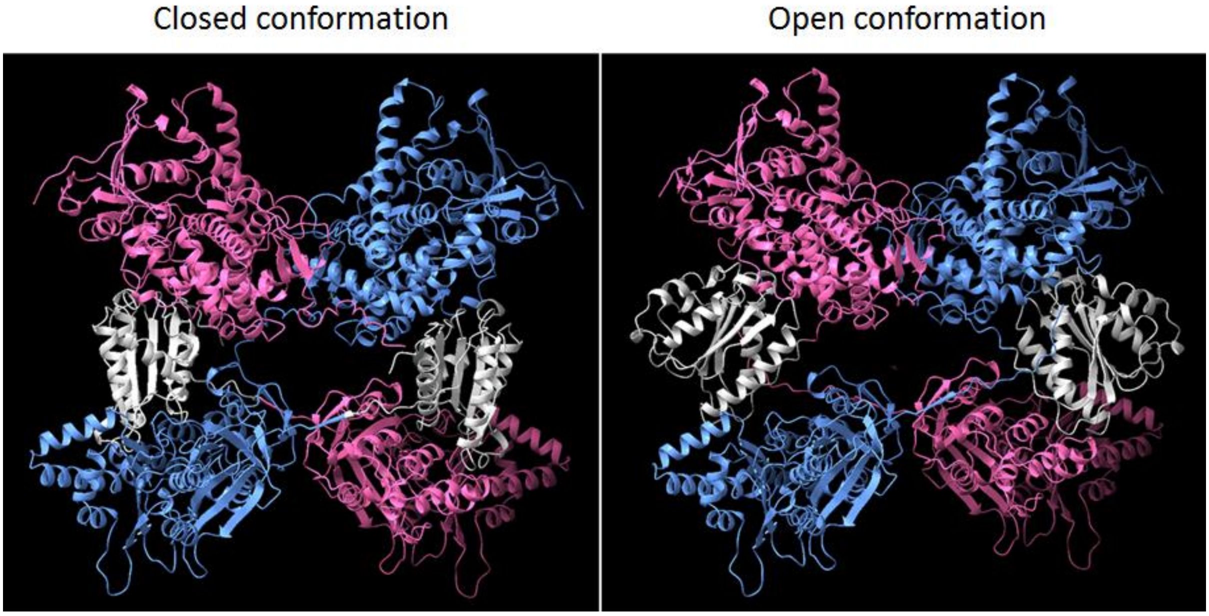

**Supplementary Figure S7.** Composite view for full length CYP102A1 hybrid dimer in which one monomer is in the closed conformation and the other monomer is in the open conformation. The model was built as described in Methods section. *Left pannel*. AF2A-based model of this hybrid dimer in a ribbon representation. *Right pannel*. Docking of the AF2A based model of the hybrid dimer in the EMD-20786 map for the open structure 1 of CYP102A1 A82F mutant[22].

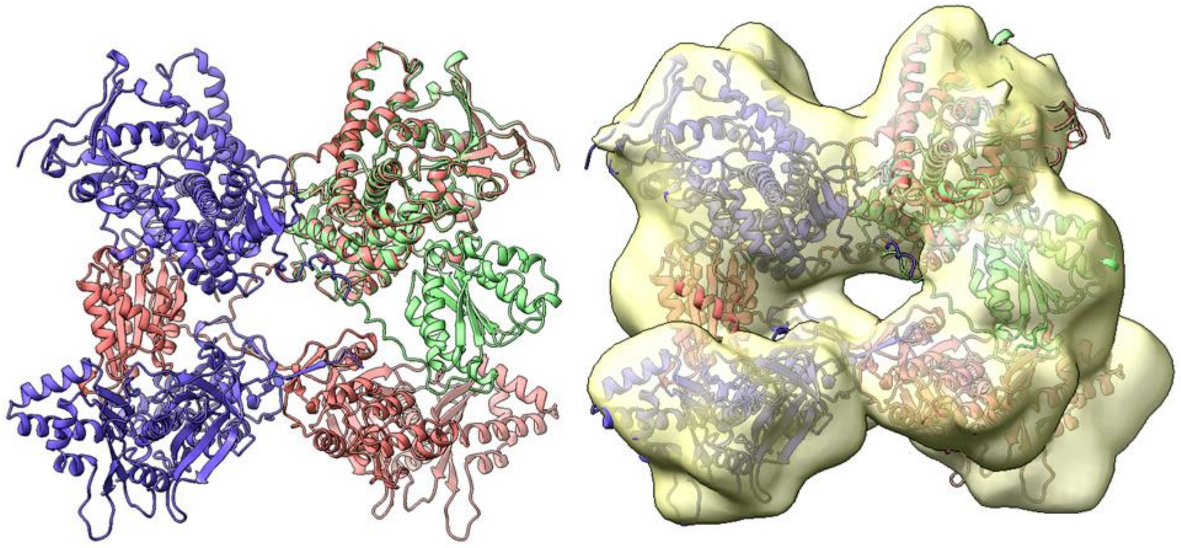

**Supplementary Figure S8.** Multiple sequence alignment showing the conserved residues at the P450d-P450d interface in CYP102A1 related enzymes. All these bacterial enzymes form a P450d-P450d dimer in AF2A models.

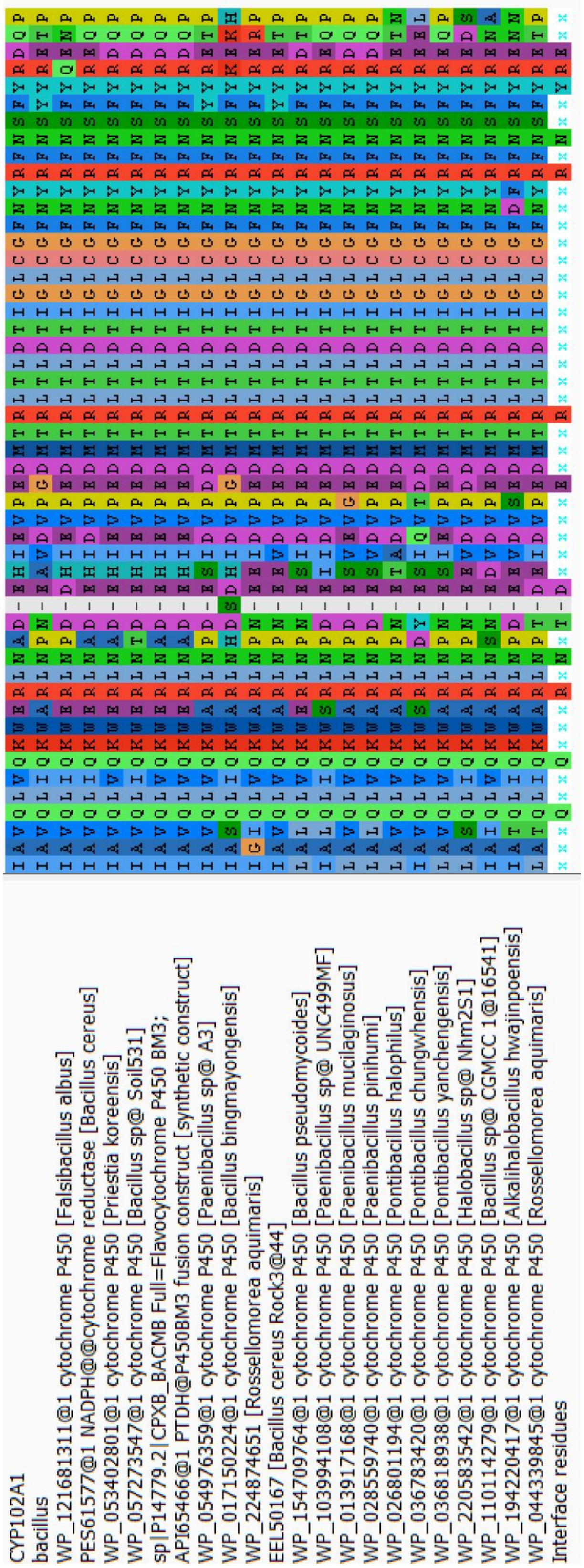

**Supplementary Figure S9.** Alternate conformations predicted for the FMNd-FADd linker region in a pair of CYP102A1 mutants. The segment $^{644}$SLQF$^{647}$ corresponding to the small β-fold at the entry of the FMNd-FADd linker in the FADd-FADd interface was changed *in silico* to the sequence $^{644}$AGAG$^{647}$, with the aim to destabilize this β-fold. Two alternate structures were randomly obtained upon repeated AF2A modeling of the same pair of this sequence variant. The structure presented in panel A was monomeric forming and intrachain FMNd-FADd complex, whereas the structure presented in panel B was dimeric forming an interchain complex similarly to the structure always found with the wild-type sequence. The FMN binding domain is colored (turquoise and pale blue) and the mutated sequence segment in red. Except for the chain connectivity (monomer *vs*. dimer), the two models were highly similar, differing only by the structure of the FMNd-FADd linker (deep blue color) that includes the mutated segment (red color). A single copy of the monomer is illustrated on panel A when the half of the dimer that includes the two-interface crossing regions was illustrated on panel B.  The red-framed boxes enlighten the corresponding sequences in model A and B: A1 with B3 and A2 with B4.

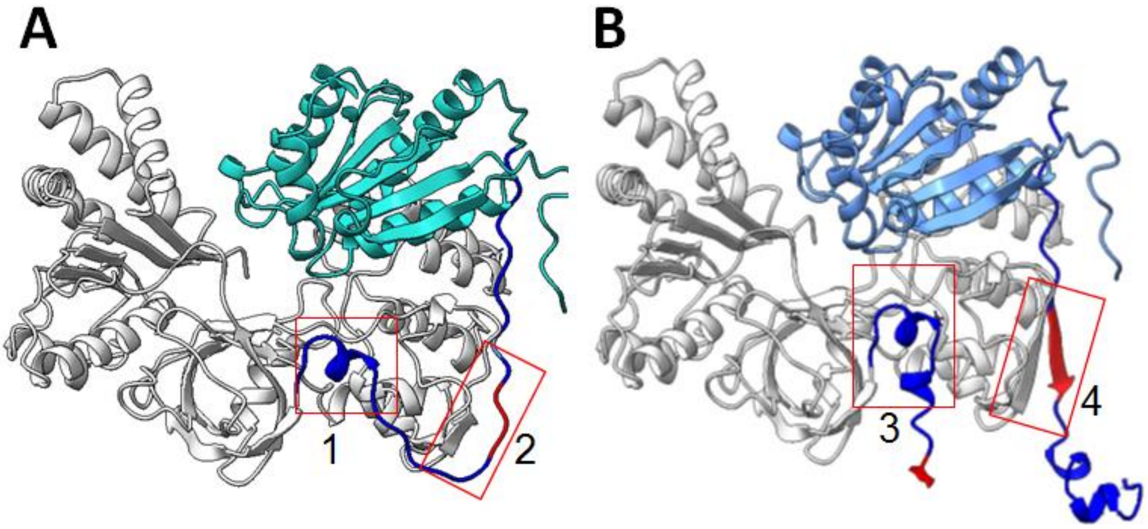

**Supplementary Figure S10.** Comparison of the AF2A predicted dimeric structures of the reductase domain found in wild-type CYP102A1 with the reductase domain of SYN001, the human-bacterial mosaic enzyme recreating the bacterial FADd-FADd interface within a eukaryotic CPR context. The structure of the mosaic enzyme was predicted to be able to dimerize, despite its amino acid sequence being mostly that of the strictly monomeric human CPR. The FMN domains are shown at the top and the FAD domains at the bottom of both structures. The two monomers of a dimer are colored differently for the sake of clarity. The structural organization of the FAD-FAD interfaces in the two enzymes is highly similar.

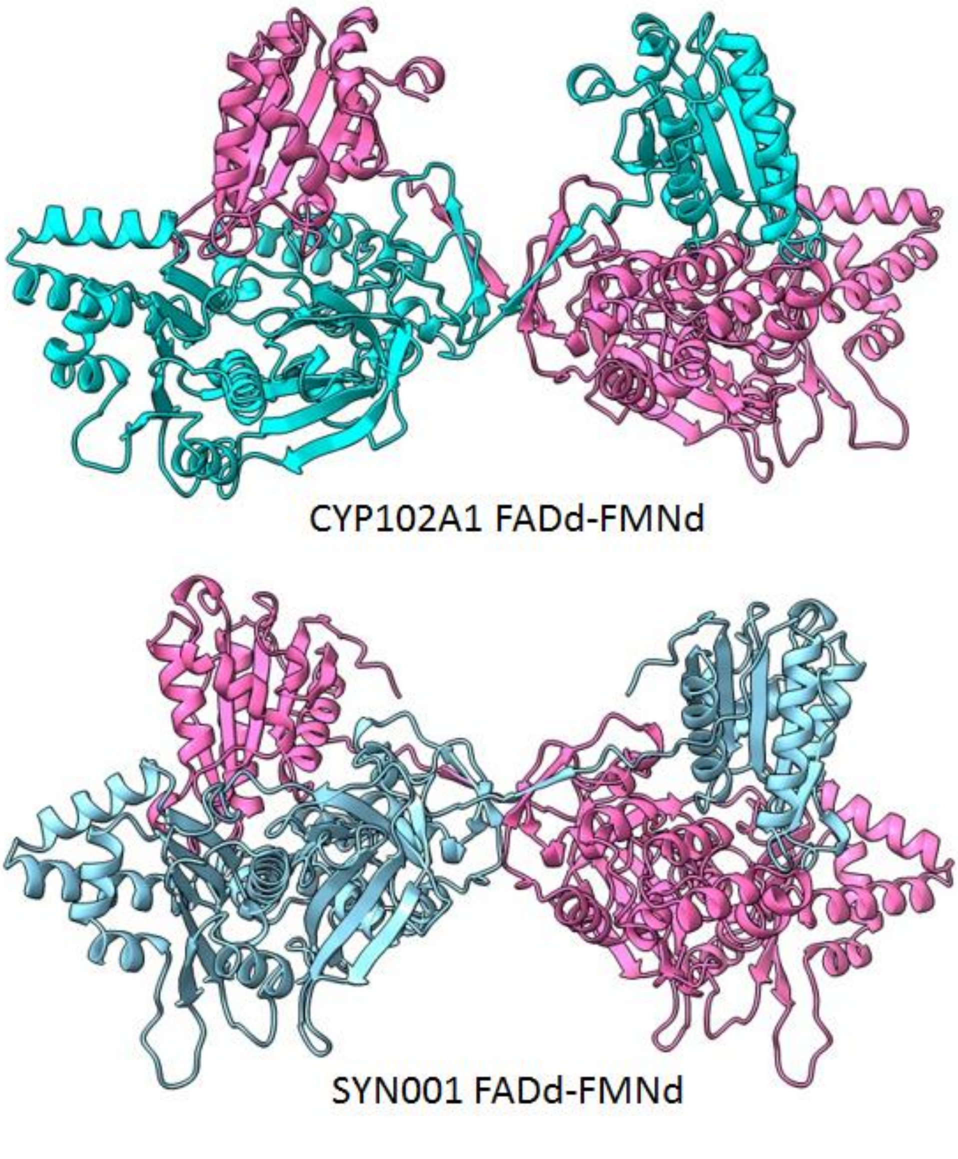

**Supplementary Figure S11.** Overlay of the structural prediction for the mosaic SYN001 reductase domain with various crystal structures. Considering the dimeric and symmetrical structure of the enzyme only a half part can be compared to monomeric CPRs. *Panel A*. Overlay of SYN001 structure (colored magenta) with the crystal structure of human CPR (PDB 3qe2) (colored blue). *Panel B*. Overlay of SYN001 structure (colored pale green) with the crystal structures of the wild-type CYP102A1 FMN domain (PDB 1bvy) and FAD domain (PDB 4dqk) (both colored magenta).

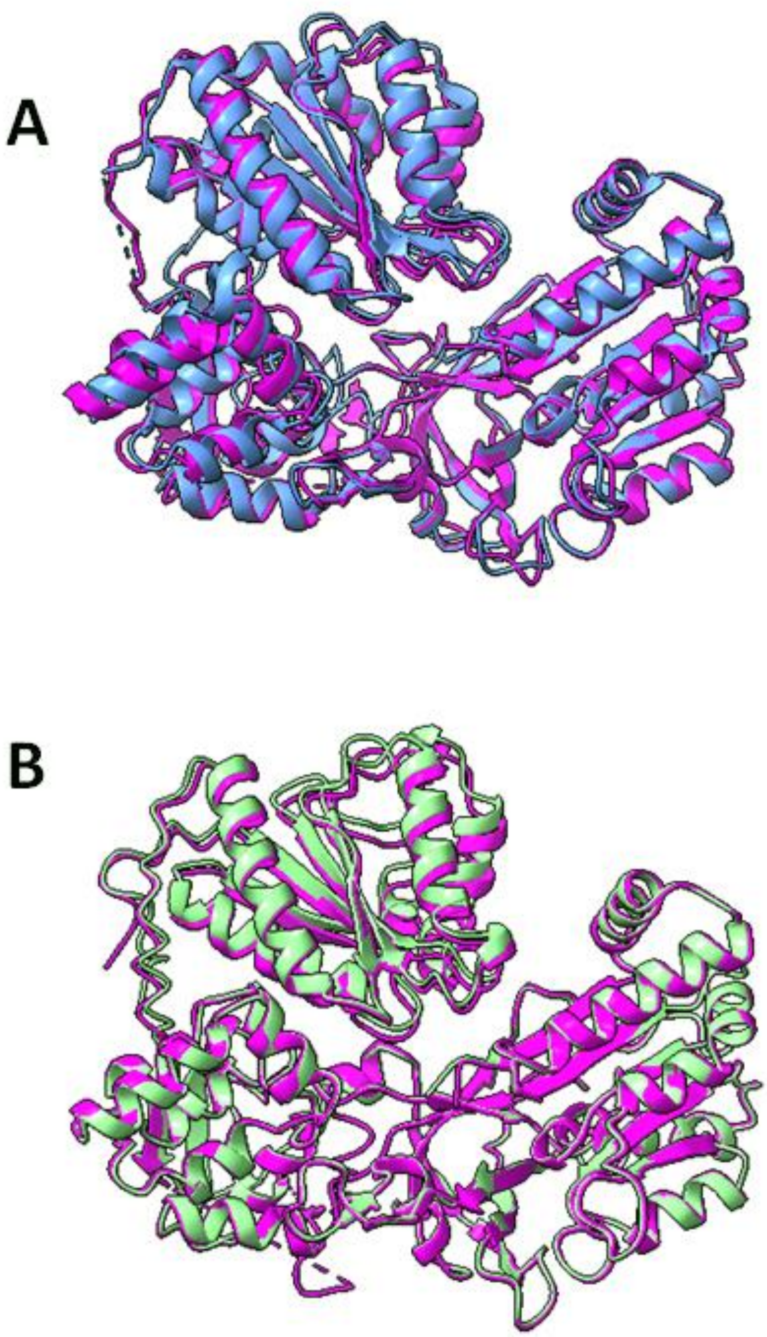

**Supplementary Table S1.** RMSD between AF2A predictions and crystallographic data for the different CYP102A1 subpart considered. Both P450d-FMNd and FMNd-FADd linker regions were excluded from calculation due to their flexibility. CYP102A1 heme domain from PDB structures 4kew and 6h1s were considered for RMSD minimization from the residue Met6 (MPQP sequence) to Lys453 (KIPL sequence). FMNd from PDB structure 1bvy was considered from the residue Asn480 (NTPLL sequence) to Tyr628 (YFNL sequence). FADd from PDB structure 4dqk was considered from His660 (in HGAF sequence) to Lys1044 (in KDVWA sequence at the C-terminus). Numbering refers to full length CYP102A1 sequence. AF2A structures used for comparisons were extracted from model of the full-length monomer for P450d and FMNd comparisons and from the closed FMNd-FADd structure for FADd comparison. Amino acid residues corresponding to unresolved parts in crystal structures were erased from AF2A models before RMSD calculations as well as heteroatoms.

| **P450d** | **Crystal structures** | |
|---|---|---|
| | 6hs1 | 4kew |
| AF2A model | 1.14 Å | 1.18 Å |
| PDB 6hs1 | - | 0.75 Å |
| **Reductase domain** | **Crystal structures** | |
| | 1bvy | 4dqk |
| AF2A FADd | - | 1.03 Å |
| AF2A FMNd | 0.88 Å | - |

**Supplementary Table S2.** Level of similarity, tendency to dimerize and domain interactions in CYP102A1 related enzymes. Predicted tendency to dimerize was evaluated as the fraction of dimeric structures predicted by AF2A for pair of P450d, and FMNd-FADd parts of the full-length enzymes. Amino acid sequence identities between aligned sequences are given as percent. The surface of the buried interface in dimers and the binding free energy of the corresponding complex association were calculated with Prodigy algorithm. Sequence id referred to GenBank entries.

| Sequence id | CYP102A1 ADB P14779 | WP_057273547 | WP_053402801 | PES61577 | WP_044339845 | COF77949 | WP_003242884 |
|---|---|---|---|---|---|---|---|
| Species | *Priestia megaterium* | *Bacillus sp* | *Priestia koreensis* | *Bacillus cereus* | *Rossellomorea aquimaris* | *Streptococcus pneumoniae* | *Bacillus subtilis* |
| P450 domain identity | 100 % | 98 % | 87 % | 79 % | 64 % | ND | 64 % |
| FMN-FAD domain identity | 100 % | 95 % | 73 % | 68 % | 57 % | 56 % | 54 % |
| Predicted as P450d dimer | 5/5 | 5/5 | 5/5 | 3/3 | 5/5 | ND | 4/5 |
| Predicted as FADd-FMNd dimer | 4/5 | 3/5 | 3/5 | 1/5 | 4/5 | 5/5 | 4/5 |
| P450d-P450d interface surface | 2473 Å$^2$ | 2713 Å$^2$ | 2536 Å$^2$ | 2309 Å$^2$ | 2377 Å$^2$ | ND | 2717 Å$^2$ |
| P450 domain ΔG binding (kcal/mol) | -9.3 | -8.7 | -10.0 | -11.5 | -10.5 | ND | -11,3 |
| FMN-FAD domain ΔG binding (kcal/mol) | -19.8 | -20.9 | -20.0 | -20.9 | -22.8 | -19.8 | -17.6 |

**Supplementary Table S3.** Sequence of modelized SYN001 mosaic enzyme. Sequence stretches from human CPR are indicated by blue letters and sequences from bacterial CYP102A1 are in red.

KMKKTGRNIIVFYGSQTGTAEEFANRLSKDAHRYGMRGMSADPEEYDLADLSSLPEIDNALV
VFCMATYGEGDPTDNAQDFYDWLQETDVDLSGVKFAVFGLGNKTYEHFNAMGKYVDKRLEQL
GAQRIFELGLGDDDGNLEEDFITWREQFWPAVCEHFGVEATGEEDNKSTLSLQFVDSANQKP
PFDAKNPFLAAVTTNRKLNQGTERHLMHLELDISDSKIRYESGDHVGVIPRNYEGIVNRVTA
RLGLDASQQIRLEAEEEKLAHLPLAKTVSVEELLQYVELQDPVTRTQLRAMAARTVCPPHKV
ELEALLEKQAYKEQVLAKRLTMLELLEKYPALRPPIDHLCELLPRLQARYYSIASSSKVHPN
SVHICAVVVEYETKAGRINKGVATNWLRAKEEGALVPMFVRKSQFRLPFKATTPVIMVGPGT
GVAPFIGFIQERAWLRQQGKEVGETLLYYGCRRSDEDYLYREELAQFHRDGALTQLNVAFSR
EQSHKVYVQHLLKQDREHLWKLIEGGAHIYVCGDARNMARDVQNTFYDIVAELGAMEHAQAV
DYIKKLMTKGRYSLDVWS

| SYN001 | Human CPR | CYP102A1 | Other residue |
|---|---|---|---|
| 1-168 | 75-242 | | |
| 169-182 | | 637-651 | |
| 183-232 | 274-323 | | |
| 233-249 | | 702-718 | |
| 250 | 341 | | |
| 251-301 | | 720-770 | |
| 302 | | | R |
| 303-341 | | 772-810 | |
| 342-403 | 440-501 | | |
| 404-405 | | 874-875 | |
| 406-576 | 510-680 | | |

**Supplementary Table S4.** Factors controlling the dimerization of CYP102A1 and related enzymes.

| Domains | Eukaryotic enzymes | Prokaryotic fusion enzymes |
|---|---|---|
| **P450** | Monomeric | Dimeric with a conserved interface |
| **Reductase** | Monomeric | Crossed dimer |
| **FMNd-FADd** | Intra-monomer complex | Inter-monomer complex |
| **FADd-FADd interface** | Non-complementary surface | Complementary surfaces with a conserved interface |
| **FMNd-FADd linker** | Long linker | Short linker |
| **4-residue β-strand at FADd-FADd interface** | Dispensable | Required |
